# A representational similarity analysis of cognitive control during color-word Stroop

**DOI:** 10.1101/2020.11.22.392704

**Authors:** Michael C. Freund, Julie M. Bugg, Todd S. Braver

## Abstract

Progress in understanding the neural bases of cognitive control has been supported by the paradigmatic color-word Stroop task, in which a target response (color name) must be selected over a more automatic, yet potentially incongruent, distractor response (word). For this paradigm, models have postulated complementary coding schemes: dorsomedial frontal cortex (DMFC) is proposed to evaluate the demand for control via incongruency-related coding, whereas dorsolateral prefrontal cortex (DLPFC) is proposed to implement control via goal and target-related coding. Yet, mapping these theorized schemes to measured neural activity within this task has been challenging. Here, we tested for these coding schemes relatively directly, by decomposing an event-related color-word Stroop task via representational similarity analysis (RSA). Three neural coding models were fit to the similarity structure of multi-voxel patterns of human fMRI activity, acquired from 65 healthy, young-adult males and females. Incongruency coding was predominant in DMFC, whereas both target and incongruency coding were present with indistinguishable strength in DLPFC. In contrast, distractor coding was strongly encoded within early visual cortex. Further, these coding schemes were differentially related to behavior: individuals with stronger DLPFC (and lateral posterior parietal cortex) target coding, but weaker DMFC incongruency coding, exhibited less behavioral Stroop interference. These results highlight the utility of the RSA framework for investigating neural mechanisms of cognitive control and point to several promising directions to extend the Stroop paradigm.

**Significant Statement:** How the human brain enables cognitive control — the ability to override behavioral habits to pursue internal goals — has been a major focus of neuroscience research. This ability has been frequently investigated by using the Stroop color-word naming task. With the Stroop as a test-bed, many theories have proposed specific neuroanatomical dissociations, in which medial and lateral frontal brain regions underlie cognitive control by encoding distinct types of information. Yet providing a direct confirmation of these claims has been challenging. Here, we demonstrate that representational similarity analysis (RSA), which estimates and models the similarity structure of brain activity patterns, can successfully establish the hypothesized functional dissociations within the Stroop task. RSA may provide a useful approach for investigating cognitive control mechanisms.

## 1 Introduction

Goals, held in mind, can be used to overcome behavioral habits. Understanding how the human brain enables such *cognitive control* has been a fundamental interest of both basic and translational cognitive neuroscience. Toward this end, the use of *response conflict* tasks has been instrumental (e.g., Botvinick, Braver, Barch, Carter, & Cohen, 2001; Ridderinkhof, Ullsperger, Crone, & Nieuwenhuis, 2004). These tasks involve trials in which a less-automatic, but goal-relevant course of action, the *target* response, must be selected in the face of a habitual, but goal-irrelevant alternative, the *distractor*. The paradigmatic example is the color-word Stroop task (MacLeod, 1991; Posner & Snyder, 1975; Stroop, 1935): on each trial, the hue of a word must be named, despite the word expressing a potentially conflicting, that is, *incongruent*, color (Figure 1, C). A major goal in this field has been to use measures of neural activity evoked by response conflict tasks, such as Stroop, to test models of cognitive control.

**Figure 1:**
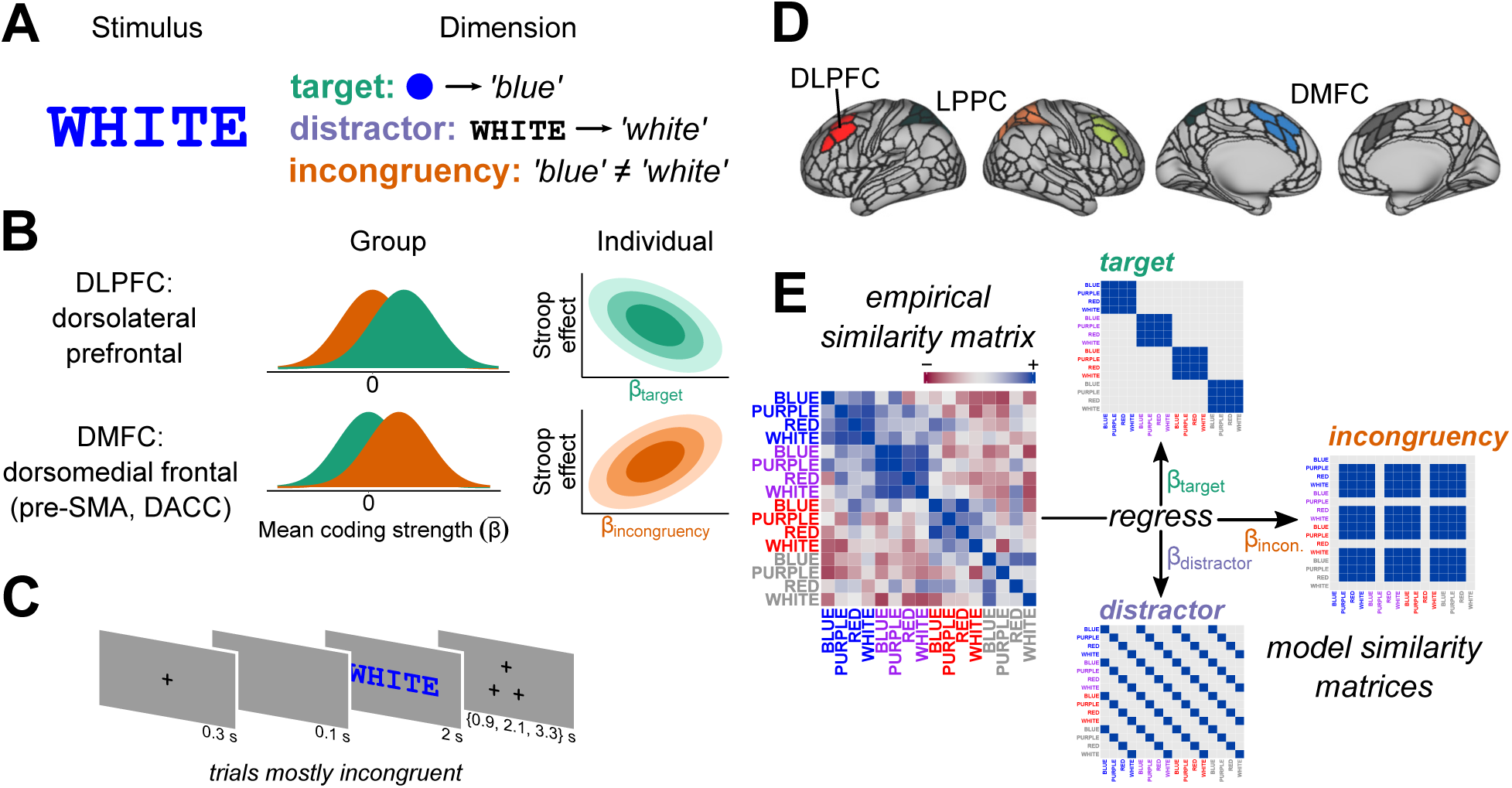
Schematic of framework and hypotheses. **A**, Conceptual framework. Cognitive theories and computational models of control have decomposed the classic color-word Stroop task into three *task dimensions*: the *target* (the goal-relevant mapping of the stimulus hue to the target response, i.e., color naming), the *distractor* (the prepotent but goal-irrelevant mapping of the stimulus word to the non-target or distractor response, i.e., word reading), and *incongruency* (whether the target and distractor responses match or mismatch). **B**, Hypotheses. Neuroscientific frameworks of cognitive control propose that representation of task dimensions is anatomically dissociated across medial and lateral frontoparietal cortices. Dorsomedial frontal cortex (DMFC), including dorsal anterior cingulate (dACC) and pre-supplementary motor area (pre-SMA), is proposed to “evaluate” demands for control, using information correlated with the *incongruency* dimension (**A**, bottom row), to signal when (and how) the current attentional or action selection policies are suboptimal (Ridderinkhof, Ullsperger, Crone, & Nieuwenhuis, 2004; Shenhav, Botvinick, & Cohen, 2013). Conversely, dorsolateral PFC (DLPFC), in concert with lateral posterior parietal cortex (LPPC), is proposed to guide, or “implement,” goal-driven attentional selection and mapping of hue–target-response processes, by way of representing information related to the goal-dependent target dimension (**A**, top row; Miller & Cohen, 2001); Buschman & Miller, 2007). Double dissociations are therefore predicted at multiple levels of analysis. At the group level, incongruency coding (orange univariate distributions) should predominate in DMFC, whereas target coding (green univariate distributions) should predominate in DLPFC. At the individual level, if the strength of target-related coding in DLPFC reflects the robustness of goal-driven selection, then subjects with stronger DLPFC target coding should resolve Stroop interference more efficiently (green bivariate distribution; Kane & Engle, 2002; Braver, 2012). Conversely, if the strength of incongruency-related coding in DMFC indicates a maladaptive selection policy, then subjects with stronger incongruency-related coding should resolve Stroop interference *less* efficiently (orange bi-variate distribution; Braver, 2012; MacDonald, Cohen, Stenger, & Carter, 2000). The *β* notation in axis titles corresponds to that used in **E** (and throughout this manuscript); 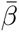 indicates mean over subjects. **C–E**, Analytic framework. Participants performed a color-word Stroop task while undergoing an fMRI scan (**C**). To derive neural correlates of the theorized task dimensions (in **A–B**), a general linear model estimated the BOLD response evoked by sixteen unique Stroop stimuli (e.g., “WHITE” displayed in blue hue) independently for each voxel. We analyzed a mostly-incongruent trial condition, in order to obtain balanced estimates of each trial type. The Glasser et al. (2016) multi-modal atlas was used to parcellate cortex (**D**, light silver borders). Contiguous sets of parcels that tiled our regions of interest (“superparcels”) were defined and treated as analytic units (see Figure 1-1 for list). Within each superparcel, linear correlations among response patterns from the sixteen stimuli were estimated to form an empirical similarity matrix (**E**, left; stimuli that were white are presented here in grey). Through rank regression, these matrices were fit to three representational models (**E**, right), which corresponded to the three hypothesized dimensions of the Stroop task (**A**). The resulting *β* coefficients summarized the extent to which a parcel emphasized, within its distributed activity patterns, the representation of each unique task dimension. These *β* coefficients were used as the primary dependent variables in group-level analyses, and primary independent variables in individual-level analyses (e.g., as in **B**). Critically, we verified the specificity of our design and analysis via simulation (Figure 1-2; *cf.*, Cai, Schuck, Pillow, & Niv, 2019).

One broad, neurocomputational-level model ascribes particular roles to different frontoparietal regions in overcoming response conflict (Miller & Cohen, 2001; Shenhav, Botvinick, & Cohen, 2013). Central to this view is the type of task information these regions encode. The dorsomedial frontal cortex (DMFC) is proposed to “evaluate” demand for cognitive control, via encoding of *incongruency-related information* (Figure 1, A, bottom row). Such information, according to this view, is used by dorsolateral prefrontal cortex (DLPFC), in concert with lateral posterior parietal cortex (LPPC), to “implement” control, via encoding of goal and *target-related information* (Figure 1, A, top row). Thus, this model predicts key functional dissociations between medial and lateral frontoparietal cortex (Figure 1, B). But, although this view has been influential, directly establishing these dissociations during the performance of standard color-word Stroop tasks has been difficult.

To date, the most traction on this problem has been gained via fMRI designs that temporally dissociate presentation of task-rule and incongruency-related information, in which subjects were instructed prior to each Stroop trial about the task to perform (color-naming, word-reading; Floden, Vallesi, & Stuss, 2010; MacDonald, Cohen, Stenger, & Carter, 2000). But, while these studies generally found supportive evidence for the key claims, results were subject to three notable limitations. First, these studies were likely underpowered for fMRI (e.g., N = 12 in Floden et al., 2010; N = 9 in MacDonald et al., 2000). This fact alone warrants a follow-up study. Second, it is unclear whether the results extend to the more-standard Stroop-task design, in which task rules are not explicitly instructed prior to each trial, but are instead internally maintained. For example, goal-relevant coding in DLPFC may depend upon such explicit rule instruction. Third, the prior results do not speak to functional dissociations *within* a single Stroop trial, during which interference is actually experienced and resolved. It is therefore possible, for instance, that the role of DLPFC (or other frontoparietal regions) in Stroop is primarily preparatory, and is less critical during actual interference resolution.

To address these questions, a neuroanatomically precise technique is needed that does not rely on temporal dissociations, but can instead read out multiple, simultaneously encoded sources of task information from individual brain regions of interest. Multivariate (multivoxel) pattern analysis (MVPA) of fMRI, in popular use for over a decade (Cox & Savoy, 2003; Edelman, Grill-Spector, Kushnir, & Malach, 1998; Haxby, Gobbini, Furey, & Ishai, 2001), accomplishes exactly this purpose.

Surprisingly, however, these methods have not been brought to bear on the question of a functional dissociation between medial and lateral frontoparietal cortex in resolving Stroop conflict.

We fill this gap in the literature by using Representational Similarity Analysis (RSA; Edelman, Grill-Spector, Kushnir, & Malach, 1998); Kriegeskorte 2008), a specific MVPA framework, to test for dissociations in frontoparietal coding during Stroop-task performance (Figure 1, C–E). We conducted a retrospective analysis of data collected as part of the Dual Mechanisms of Cognitive Control project (Braver, Kizhner, Tang, Freund, & Etzel, 2020). Our primary goal was a proof-of-principle: to demonstrate the potential of RSA for testing theorized distinctions in neural coding within cognitive control tasks, such as the Stroop (Freund, Etzel, & Braver, 2021).

## 2 Materials and Methods

We report how we determined our sample size, all data exclusions, all manipulations, and all measures in the study (Simmons, Nelson, & Simonsohn, 2011).

### 2.1 Code, Data, and Task Accessibility

Code (R Core Team, 2019) and data to reproduce all analyses, in addition to supplementary analysis reports, are publicly available (https://doi.org/10.5281/zenodo.4784067). As part of the planned data release of the Dual Mechanisms of Cognitive Control project, raw and minimally preprocessed fMRI data have been deposited on OpenNeuro (10.18112/open-neuro.ds003465.v1.0.3). Additionally, the authors will directly share the specific fMRI data used for this study upon reasonable request. Task scripts are available at the DMC project website (http://pages.wustl.edu/dualmechanisms/tasks). More detailed information regarding all aspects of the project can be found on the Project’s OSF page (https://osf.io/xfe32/).

### 2.2 Participants

Individuals were recruited from the Washington University and surrounding St Louis metropolitan communities for participation in the Dual Mechanisms of Cognitive Control project. The present study began with a subset (*N* = 66; 38 women, 26 men, 1 “prefer not to answer”) of these subjects: those with a full set of imaging and behavioral data from the Stroop task during a particular scanning session (the *proactive* session), selected for methodological reasons (see *Selection of Data*). One subject was excluded from all analyses due to a scanner error. We split the remaining sample into two sets of individuals: a *primary analysis set* (*N* = 49; 27 women, 21 men, 1 “prefer not to answer”), which we used in all analyses, and a *validation set* (*N* = 16; 11 women, 5 men), which was only used in the *Model Selection* analysis (see below). This unbalanced partitioning was done to account for the familial structure present within our sample. Specifically, subjects within each set (primary analysis, validation) were all unrelated; however, subjects within the validation set were co-twins of 16 subjects within the primary analysis set. Two of these co-twins were selected for use in the *primary analysis set* as their respective co-twins had atypically high rates of response omission (*>*10%; *>*20% errors of any type); the remaining co-twins were randomly selected. Critically, partitioning the sample in this way ensured that the primary analysis set was a random sample of independent subjects.

The partitioning of the data into two subsets also afforded the opportunity to use the validation subset as held-out data for evaluation of the brain–behavior model within the *Model Selection* analysis. As we performed this sorting of individuals into primary and validation sets only once and did not analyze the validation-set data except to assess predictive accuracy of the final selected model, the validation set provides an unbiased assessment of predictive accuracy, in the sense that no statistical “double-dipping” could have occurred. But because the sets are familially dependent, it is perhaps more accurate to consider the validation-set analyses as assessing a kind of test–retest reliability (i.e., while eliminating the potential confound of practice effects), rather than providing an estimate of out-of-sample predictive accuracy. To evaluate this matter, follow-up control analyses were conducted in which the co-twins were removed from the primary analysis set.

### 2.3 Experimental Design and Statistical Analysis

#### 2.3.1 Task

Participants performed the verbal color-word Stroop (1935) task. Names of colors were visually displayed in various hues, and participants were instructed to “say the name of the color, as fast and accurately as possible; do not read the word”.

The set of stimuli consisted of two intermixed subsets of color-word stimuli (randomly intermixed during the task): a *mostly incongruent* and an *unbiased* set. Each stimulus set was created by pairing four color words with four corresponding hues in a balanced factorial design, forming 16 unique color-word stimuli within each set. The *mostly incongruent* set consisted of stimuli with hues (and corresponding words) ‘blue’ (RGB = 0, 0, 255), ‘red’ (255, 0, 0), ‘purple’ (128, 0, 128), and ‘white’ (255, 255, 255); the *unbiased* group, of ‘black’ (0, 0, 0), ‘green’ (0, 128, 0), ‘pink’ (255, 105, 180), and ‘yellow’ (255, 255, 0). These words were centrally presented in uppercase, bold Courier New font on a grey background (RGB = 191, 191, 191). Of stimuli within the *mostly incongruent* set, incongruent stimuli were presented to subjects more often than congruent stimuli (per block, proportion congruent = 0.25). *Unbiased* stimuli were presented with a balanced frequency (proportion congruent = 0.5). These manipulations of incongruency statistics are standard manipulations to elicit proactive control (Bugg, 2014; e.g., Gonthier, Braver, & Bugg, 2016) and were performed to investigate questions outside the scope of the current study. Thus, as described further below, the unbiased stimulus set was excluded from all analyses.

Each trial (e.g., Figure 1, C) began with a central fixation cross, presented for 300 ms on a grey background (RGB = 191, 191, 191). The color-word stimulus, preceded by a blank screen following fixation offset (100 ms), was centrally presented for a duration of 2000 ms, fixed across trials. The duration of the inter-trial interval (triangle of fixation crosses) was either 900, 2100, or 3300 ms, selected randomly (with uniform probability). Each of two scanning runs consisted of three blocks of 36 trials, intermixed with 4 resting fixation blocks, during which a fixation cross appeared for 30 s. This formed a mixed block-event design (Petersen & Dubis, 2012). Each of the 16 *mostly incongruent* stimuli — that is, each unique colored word (e.g., “BLUE” displayed in red hue) — were presented in both runs. Within each run for each participant, *mostly incongruent* stimuli were presented an equal number of times within each block. Within each block, stimulus order was fully randomized.

#### 2.3.2 Selection of Data

We focused our fMRI pattern analyses solely on trials from the *mostly incongruent* stimulus set within a particular scanning session (the “proactive” session) of our Stroop task. This selection was made purely on the basis of methodological reasoning: these trials were the only set of trials within the larger Dual Mechanisms project in which each unique Stroop stimulus (i.e., one of the 16 color-word combinations) was presented an equal number of times (9) to each participant, constituting a balanced design. Balanced designs ensure that differences in the total number of trials per condition cannot explain any differences observed in pattern correlations among conditions.

#### 2.3.3 Display and Recording Systems

The experiment was programmed in EPrime 2.0 (“E-Prime 2.0,” 2013), presented on a Windows 7 Desktop, and back-projected to a screen at the end of the bore for viewing via a mirror head-mount. Verbal responses were recorded for offline transcription and response-time (RT) estimation. The first 45 participants spoke into a MicroOptics MR-compatible electronic microphone (MicroOptics Technologies, Inc.); due to mechanical failure, however, we replaced this microphone with the noise-cancelling FOMRI III (OptoAcoustics, Ltd.), which subsequent participants used. A voice-onset processing script (from the MATLAB Audio Analysis Library) was used to derive response-time estimates on each trial via spectral decomposition (the accuracy of which was verified by manually coding response times from a subsample of subjects and ensuring the two methods gave similar estimates). Code for this algorithm is available within the Dual Mechanisms GitHub repository (https://github.com/ccplabwustl/dualmechanisms/tree/master/preparationsAndConversions/audio).

Importantly, we verified that the change in microphone did not induce confounding between-subject variance in response-time measures of interest. While response time estimates recorded via the Micro-Optics microphone tended to be slower (*b* = 102.59*, p* = 0.01) and more variable (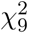 = 3655*, p* = 0), the magnitude of the Stroop effect was not observably impacted by the microphone change (*b* = −5.88*, p* = 0.64).

#### 2.3.4 Image Acquisition, Preprocessing, and Generalized Linear Model

The fMRI data were acquired with a 3T Siemens Prisma (32 channel head-coil; CMRR multi-band sequence, factor = 4; 2.4 mm isotropic voxel, with 1200 ms TR, no GRAPPA, ipat = 0), and subjected to the minimally pre-processed functional pipeline of the Human Connectome Project (version 3.17.0), outlined in (Glasser et al., 2013). More detailed information regarding acquisition can be found on the Project OSF site (https://osf.io/tbhfg/). All analyses were conducted in volumetric space; surface maps are displayed in figures only for ease of visualization. Note that prior to revision of this manuscript, the data were re-processed with fMRIPrep, using the standard fMRIPrep pipelines (Esteban et al. 2019; 2018; see https://fmriprep.org/en/stable/citing.html#note-for-reviewers-and-editors for more information). At this point, the pre-processed results with the HCP pipelines were inadvertently removed. Thus, some follow-up control analyses were conducted with the fMRIPrep-preprocessed data (Figure 2-4, Figure 3-2, Figure 4-4). The fMRIPrep pipeline was implemented in a Singularity container (Kurtzer, Sochat, & Bauer, 2017) with additional custom scripts used to implement file management (more detail on the pipeline is available at https://osf.io/6p3en/; container scripts are available at https://hub.docker.com/u/ccplabwustl).

After pre-processing, to estimate activation patterns, we fit a whole-brain voxel-wise generalized linear model (GLM) to blood-oxygen-level dependent (BOLD) time-series in AFNI, version 17.0.00 (R. W. Cox, 1996). To build regressors of primary interest, we convolved with a hemodynamic response function [via AFNI’s *BLOCK(1,1)*] 16 boxcar timecourses, each coding for the initial second of presentation of a *mostly incongruent* stimulus that resulted in a correct response. We also included two regressors [similarly created via *BLOCK(1,1)*] to capture signal associated with congruent and incongruent trials of non-interest (*unbiased* stimuli) that prompted correct responses, an “error regressor” coding for any trial in which a response was incorrect or omitted (via *BLOCK*), a sustained regressor coding for task versus rest (via *BLOCK*), a transient regressor coding for task-block onsets [as a set of piecewise linear spline functions via *TENTzero(0,16.8,8)*], six orthogonal motion regressors, five polynomial drift regressors (order set automatically) for each run, and an intercept for each run. These models were created via *3dDeconvolve* and solved via *3dREMLfit*. The data for each subject’s model consisted of 2 runs *×* 3 blocks *×* 36 trials (144 from the *mostly incongruent* stimulus group, 72 from *unbiased*). Frames with FD *>* 0.9 were censored.

#### 2.3.5 Definition of Regions of Interest

Our primary hypotheses concerned a set of six anatomical regions: DMFC, DLPFC, and LPPC in each hemisphere. Consequently, our primary analyses employed a targeted ROI-based analysis approach. Rather than defining functional regions of interest (ROIs) via a whole-brain searchlight, which has known issues (Etzel, Zacks, & Braver, 2013), we defined ROIs via a cortical parcellation atlas. We selected the MMP atlas (Glasser et al., 2016) for two reasons. (1) The atlas was developed recently via multi-modal imaging measures, and (2) individual MMP parcels are relatively interpretable, as they are heterogeneously sized and have been explicitly connected to a battery of cognitive tasks (Assem, Glasser, Van Essen, & Duncan, 2020), the canonical functional connectivity networks (Ji et al., 2019), and a large body of neuroanatomical research (Glasser et al., 2016). We used a volumetric version, obtained from https://figshare.com/articles/HCP-MMP1_0_projected_on_MNI2009a_GM_volumetric_in_NIfTI_format/3501911?file=5534024 (also available on the project GitHub repository; see *Code, Data, and Task Accessibility*). We then defined a set of six spatially contiguous sets of MMP parcels (three in each hemisphere), which we refer to as “superparcels,” that corresponded to each of our ROIs. For full superparcel definitions, see Figure 1-1. “DMFC” was defined as the four parcels covering SMA–pre-SMA and DACC. “DLPFC” was defined as the four parcels that cover middle frontal gyrus (i.e., mid-DLPFC). “LPPC” was defined as all parcels tiling IPS (both banks and fundus), from the occipital lobe to primary somatosensory cortex. The overwhelming majority of parcels that met these anatomical criteria were assigned, within a previous report, to the cinguloopercular (most of DMFC), frontoparietal (most of DLPFC), and dorsal-attention (most of LPPC) control networks (Ji et al., 2019). Further, these ROI definitions contain several parcels that correspond to key nodes within the “multiple demand” network (Assem, Glasser, Van Essen, & Duncan, 2020). To assess the robustness of our results to particular superparcel definitions, we additionally used alternative, more inclusive, superparcel definitions of DMFC and DLPFC (see Figure 1-1). For the brain–behavior model selection analysis (see *Model Selection*), we compiled a larger set of anatomically clustered MMP parcels, covering regions across the cortex (Figure 3-3). Two additional, non-MMP ROIs were included in this set, in order to give better coverage of particular functional brain regions. A mask for ventral somatomotor cortex (the “SomatoMotor–Mouth” network) was obtained from the Gordon atlas (Gordon et al., 2016), as the MMP does not split somatomotor cortex into dorsal and ventral divisions. A mask for left ventral occipito-temporal cortex (encompassing the “visual word-form” area), was obtained using MNI coordinates −54 < *x* < −30, −70 < *y* < −45, −30 < *z* < −4, specified in a prior report (Twomey, Kawabata Duncan, Price, & Devlin, 2011). To remove cerebellar voxels from this ROI, we used the Deidricshen atlas (Diedrichsen, 2006) hosted by AFNI (https://afni.nimh.nih.gov/pub/dist/atlases/SUIT_Cerebellum/SUIT_2.6_1/).

#### 2.3.6 Estimation of Coding Strength *β*

To estimate the regional strength of target, distractor, and incongruency coding, we used the representational similarity analysis framework (RSA; Kriegeskorte, 2008). The RSA framework consists of modeling the observed similarity structure of activation patterns with a set of theoretically specified model similarity structures (Figure 1, E). For a given subject and cortical region, fMRI GLM coefficient estimates for each of the 16 conditions of interest (4 colors factorially paired with 4 words; e.g., the word “WHITE” presented in blue hue) were assembled into a condition-by-voxel activity pattern matrix **B**. The observed similarity structure was estimated as the condition-by-condition correlation matrix **R** = Cor(**B**). Cell *R_ij_* of this matrix gives the linear correlation observed between activity patterns evoked by conditions *i* and *j*. Model similarity structures were specified in this same correlation matrix form. The target model assumed that conditions (stimuli) with the same hue will evoke identical patterns, regardless of whether the words or congruency match (or mismatch). That is, if the hue of condition *i* = *j*, this model predicts *R_ij_* = 1, otherwise 0. (In the ‘target’ matrix in Figure 1, E, the only cells equal to 1 — i.e., blue cells — are those in which the stimulus hues match.) The distractor model assumed that conditions with the same word will evoke identical patterns, regardless of the hue or congruency. (If the word of condition *i* = *j*, *R_ij_* = 1, otherwise 0. In the ‘distractor’ matrix in Figure 1, E, the only cells equal to 1 are those in which the stimulus words match.) The incongruency model assumed that only conditions that were incongruent would evoke identical patterns, regardless of the hue or word. (If *i* and *j* are both incongruent, *R_ij_* = 1, otherwise 0. In the ‘incongruency’ matrix in Figure 1, E, the only cells equal to 1 are those in which both stimuli are incongruent.) As a covariate of non-interest, we also included a model capturing similarity between congruent conditions (if *i* and *j* are both congruent, *R_ij_* = 1, otherwise 0). We additionally examined alternative approaches to RSA modeling of incongruency, to see if our results were robust to this parameterization of incongruency and congruency (see Method section *Alternative RSA Incongruency Models*).

These four models were jointly fitted to the observed similarity structure from each region through multiple regression (ordinary least-squares), separately for each subject. The response vector **y** and design matrix of this regression were assembled in a series of steps. (1) The 120 unique off-diagonal elements of of each similarity matrix (one observed and four models) were extracted and unwrapped into vectors. (2) The four model similarity vectors were separately z-scored and assembled into columns. This formed the RSA design matrix. (3) The observed similarity vector was rank-transformed (Nili et al., 2014) then z-score standardized, to form a vector **r**. (4) The vector **r** was prewhitened to remove a specific nuisance component. This component stemmed from the task design: though each *mostly incongruent* stimulus occurred an equal number of times throughout the course of a session, these stimuli were not fully balanced across the two scanning runs. Specifically, half of the stimuli were presented three times in the first run versus six in the second (vice versa for the other half). As each scanning run contains a large amount of run-specific noise (Alink, Walther, Krugliak, Bosch, & Kriegeskorte, 2015; Mumford, Davis, & Poldrack, 2014), this imbalance across runs could lead to a bias in the resulting *β* coefficients, in which pattern similarity of stimuli that mostly occurred within the same run would be inflated. We formalized this component of bias as another model similarity vector, **v**, with elements equal to 1 if *the run in which condition i most frequently occurred* = *the run in which condition j most frequently occurred*, otherwise 0. The magnitude of this bias was estimated as the slope term *b*_1_ in a linear regression **r** = **v***b*_1_ + *b*_0_ + ***E***, where *b*_0_ is the intercept coefficient and ***E*** is the residual vector. The model **v** was scaled by its magnitude then subtracted from **r**, forming the RSA response vector **y** = **r** − **v***b*_1_. We additionally used an alternative, downsampling technique, to verify that our primary findings were robust to this issue (see Method section *Downsampling Analysis*).

Thus the RSA regression yielded three *β* coefficients of interest: *β*_target_, *β*_distr._, *β*_incon._. These coefficients can be understood as a (standardized) contrast on (rank-transformed) correlations of activity patterns, between conditions in which only one task dimension was shared (e.g., the target dimension for *β*_target_), versus those in which no dimensions were shared (i.e., different levels of target, distractor, and congruency).

#### 2.3.7 Dimensionality Reduction

We used non-metric multidimensional scaling (Kruskal, 1964), a flexible, non-parametric dimensionality reduction technique, to visualize the structures of activity patterns within selected regions (Figure 2): ventral somatomotor cortex (corresponding to ‘mouth’ homunculi), primary visual cortex (V1), and our (left) DMFC superparcel. These parcels were selected to highlight coding of each task dimension. For each selected region, we averaged observed correlation matrices across subjects, then subtracted these values from 1 to obtain a dissimilarity matrix. Before averaging, we *z*-transformed (inverse hyperbolic tangent, artanh) correlations, and inverted this transform after averaging. (In contrast to the RSA regression above, we did not rank-transform correlation matrices, as non-metric multidimensional scaling incorporates a monotonic regression). Similar to our RSA, we prewhitened each similarity matrix prior to conducting this procedure (see Step 4 in *Estimation of Coding Strength β*). Each mean dissimilarity matrix was submitted to an implementation of Kruskal’s non-metric multidimensional scaling, *vegan::metaMDS()* in R, to generate a 2-dimensional configuration (Oksanen et al., 2019).

**Figure 2:**
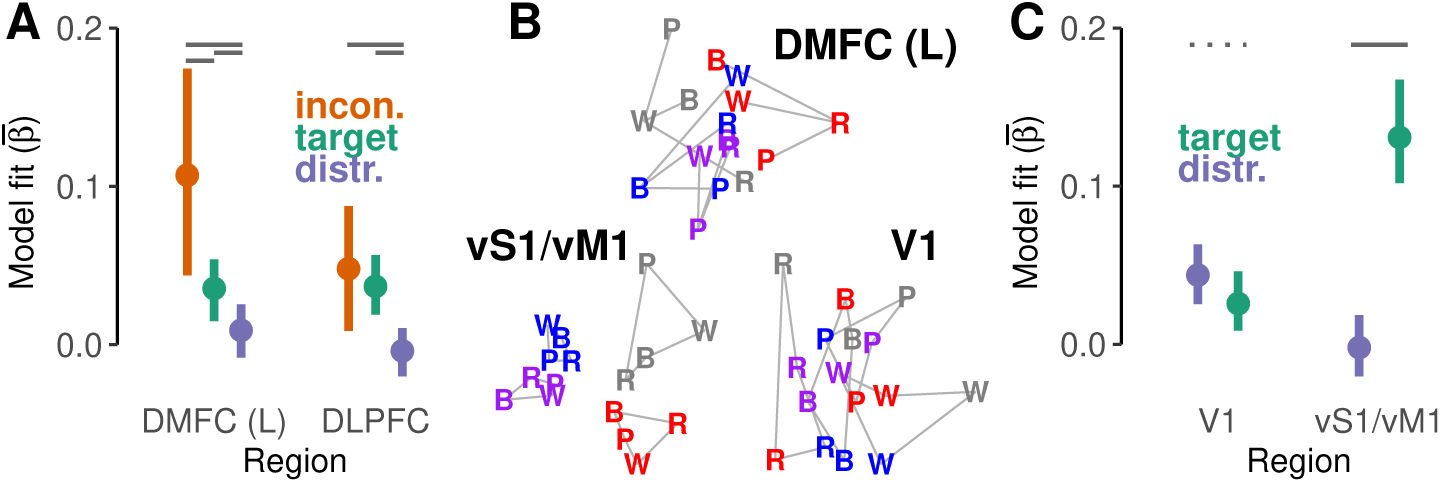
Group-level results. **A**, A single dissociation between dorsolateral versus dorsomedial frontal cortex (DLPFC, DMFC). For simplicity of display, DLPFC estimates are averaged across hemisphere (see Table 2 for per-hemisphere means for each region). Error bars show 95% confidence intervals of between-subject variability (estimated via bias-corrected and accelerated bootstrap). Horizontal grey lines at the top indicate significance of within-subject significance tests (see Table 2 for contrast estimates). While all ROIs encoded target and incongruency information, and did so more strongly than distractor information (coding of which was not detected), incongruency coding was stronger in left DMFC versus DLPFC. These results were generally robust to different analysis decisions and implementations (Figures 2-1, 2-2, 2-4, 2-5). Further, a univariate analysis failed to detect significantly higher activation on incongruent versus congruent trials in these regions (Figure 2-3). **B**, Across-voxel activity patterns from three select regions, embedded within a two-dimensional space (via non-metric multidimensional scaling). These geometries exemplify regional dissocations in coding of Stroop-task dimensions. Letters represent the first letter of the corresponding distractor word (BLUE, PURPLE, RED, WHITE), and hues represent the target color (stimuli that were white are presented here in grey). Connecting lines are drawn to highlight the various structures present. The geometry of (left) dorsomedial frontal cortex (DMFC) is marked by a radial separation of patterns evoked by congruent and incongruent conditions (lines connect congruent and incongruent stimuli of each target color), such that incongruent patterns tend to be located more centrally (i.e., more similar to each other), whereas congruent patterns peripherally diverge (i.e., less similar to each other). This suggests that incongruent trials drove a common component of activation in this region, regardless of target or distractor features. In contrast, within primary ventral somatomotor cortices (vS1/vM1), patterns strongly cluster by target level (color), whereas in primary visual cortex (V1), they tend to cluster by distractor level (word). This pattern formed a double dissociation (**C**), and indicated our distractor model was adequately powered to detect distractor coding in sensory regions (*cf.*, **A**). **C**, Plotting conventions follow those in **A**; horizontal dotted lines indicate 0.1 *> p >* 0.05 (see text for estimates).

#### 2.3.8 Group-Level Dissociation Analysis

To test for regional dissociations in coding preferences, we fit a hierarchical linear model on RSA model fits (see Estimation of Coding Strength *β*) obtained from our three ROIs within each hemisphere, and for our three RSA models. Fixed effects were estimated for the interaction of RSA model, ROI, and hemisphere. Random effects by subject were estimated for the interaction of RSA model and ROI, with a full covariance structure (9 *×* 9; Barr, Levy, Scheepers, & Tily, 2013). This model was fit with *lme4::lmer()* in R (Bates, Maechler, Bolker, & Walker, 2014).

Planned contrasts on the fixed effects were performed to test our hypotheses. P-values were estimated using an asymptotic *z*-test, as implemented by the *multcomp::glht()* function in R (Hothorn, Bretz, & Westfall, 2008). We performed three types of contrasts: (1) to compare coding strengths within-region (e.g., DMFC: incon.−target), (2) to compare between regions, within-model (e.g., target: DLPFC − DMFC), and (3) to test their interaction [e.g., (target − incon.) · (DLPFC − DMFC)]. These contrasts were performed first by collapsing across hemisphere, then within each hemisphere separately. As we did not have any hypotheses regarding lateralization, *p*-values from hemisphere-specific contrasts were FDR corrected across hemispheres.

#### 2.3.9 Selection of Behavioral Measures for Individual-Level Analyses

Audio recordings of verbal responses were transcribed and coded for errors offline by two researchers independently. Discrepancies in coding were resolved by a third. Errors were defined as any non-target color word spoken by a subject prior to utterance of the correct response (e.g., including distractor responses, but not disfluencies) or as a response omission. Trials in which responses were present but unintelligible (e.g., due to high scanner noise or poor enunciation) were coded as such.

We fit two hierarchical models on these data, one on errors and one on response times (RT). Several observations were excluded from these models. From the error model, only trials with responses coded as “unintelligible” were excluded (54). From the RT model, several types of trials were excluded: Trials with RTs greater than 3000 ms (1) or less than 250 ms (53; 52 of which were equal to zero). A cluster of fast and unrealistically invariable RTs from two subjects (23/216, 26/216) that were likely due to an artifact of insufficient voice-onset signal within the recording. Trials with a residual RT that was more extreme than 3 inter-quartile ranges from an initial multi-level model fitted to all subjects data (of the structure of the model equation below; *a la* Baayen & Milin, 2010). All trials with incorrect (137), unintelligible (54), or no response (52). In total, 232 trials were excluded (0–62 per subject), leaving 10352 trials for analysis (154–216 per subject).

RTs for subject *s* were modeled (following Laird & Ware, 1982 notation):

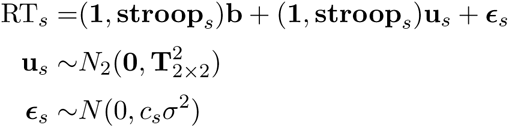

where RT*_s_* is a column vector of the response times from subject *s*, **1** is an all-ones vector (intercepts), **stroop***_s_* is a vector indicating incongruent trials, and *c_s_* is a subject-specific parameter by which their residual variance was scaled. Critically, **b** and **u***_s_* contained coefficients corresponding to the classical Stroop interference effect contrast (incongruent – congruent), for the group (**b**) and subject (**u***_s_*) levels. The scaling parameters *c_s_* relaxed the assumption that each subject had a common residual variance; a well-warranted complexity, given the vastly improved fit of the heterogeneous-variance model (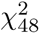*, p* = 0, ΔBIC = −3856, ΔAIC = −4203). To accomplish this, the RT model was fitted with *nlme::lme()* in R (Pinheiro, Bates, DebRoy, Sarkar, & R Core Team, 2019). The behavioral error model had similar predictors, but assumed binomially distributed error, with logit link function (fitted in *lme4*).

This hierarchical modeling framework enabled us to to estimate the amount and internal consistency of individual variability in the Stroop interference effect within both RTs and errors while accounting for trial-level error (e.g., Haines et al., 2020). We used these subject-level estimates to validate that our behavioral measures met prerequisite properties for individual differences analyses. To assess the amount of between-subject variability in the Stroop interference effect, a nested model comparison was conducted, in which the models fitted above were compared to a random-intercept model. Stroop effects differed significantly across subjects in RT (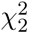 = 87.53*, p* = 0.00; ΔBIC = −69.08; ΔAIC = −83.53), but not in accuracy (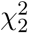 = 0.69*, p* = 0.71; ΔBIC = 18; ΔAIC = 3). We therefore did not further analyze accuracy. To assess internal consistency (defined here as cross-run correlation), we fit a model with separate congruency factors (fixed and random) per run, and a full 4 *×* 4 covariance structure, which was used to obtain the cross-run correlation in Stroop effect. Individuals’ Stroop interference effects in RT were estimated to be highly consistent across scanning runs (*r* = 0.95). We therefore considered our measurements of RT to be adequate for individual differences analysis.

#### 2.3.10 Individual-Level Dissociation Analysis

Similar to our *Group-Level Dissociation Analysis*, we tested our individual-level hypotheses within a hierarchical modeling framework. Preliminary analyses suggested error measures were inadequate for individual differences analyses (see *Materials and Methods*, *Selection of Behavioral Measures for Individual-Level Analyses*), so we focused solely on RT measures.

We began with the RT model described in the preceding section. However now, for a given RSA model and region of interest, we incorporated into the fixed effects each subject’s estimated coding strength, *β_s_*, by interacting this coding-strength term with the congruency factor. This formed a cross-level, continuous-by-categorical interaction, *β_s_* · **stroop***_s_*. The coefficient on this interaction term described how the Stroop interference effect (within-subject) varied across subjects as a function of their coding strength (between-subject). To test our hypotheses, we performed contrasts on these interaction coefficients, which are outlined within the *Results* describing Figure 3.

**Figure 3:**
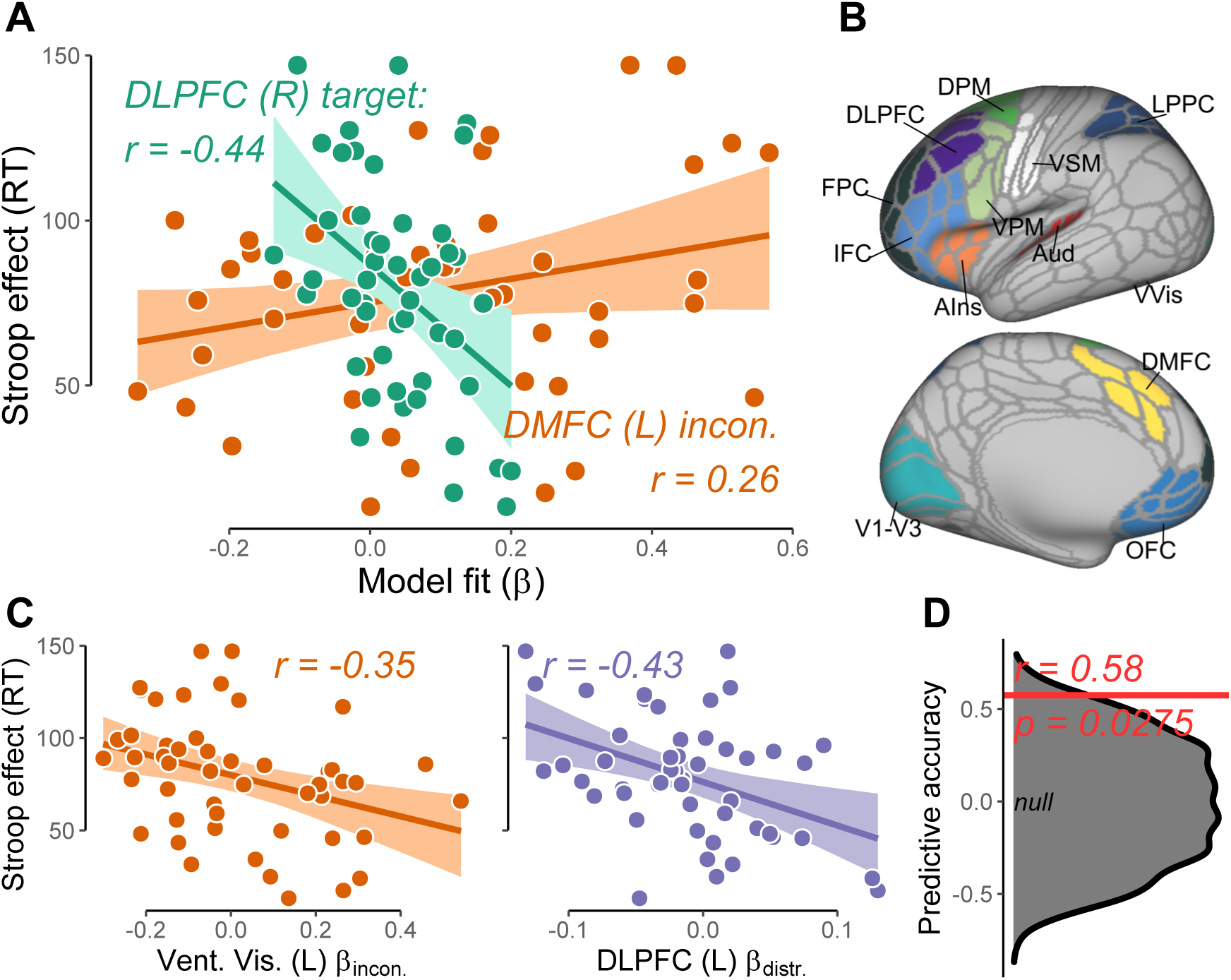
Individual-level results. **A**, The coding strength of task variables (*β*) within right dorsolateral prefrontal cortex (DLPFC) and left dorsomedial frontal cortex (DMFC) accounted for individual differences in the size of the Stroop interference effect (response time, RT) in hypothesized ways. Inset text displays linear correlation coefficients (*r*) and 95% confidence intervals (estimated via bias-corrected and accelerated bootstrap). Lines indicate a regression line of RT onto Model Fit (*β*), and confidence bands indicate 95% confidence intervals of predicted values from this regression obtained via percentile bootstrap. The direction of these observed brain– behavior relationships were robust to alternative, more conservative RSA estimation techniques (cross-run RSA and cross-validated RSA; Figure 3-2). For display of relationships between target and incongruency coding in all ROIs (DLPFC, DMFC, LPPC) and Stroop effect, see Figure 3-1. For contrasts on slopes (interaction tests between ROI and RSA model), see Table 3. **B**, A larger set of 24 cortical regions were defined for use in a data-driven model selection analysis. Surface maps display in various colors all left-hemisphere regions (excluding left lateral occipito-temporal cortex); see Figure 2-1 for full list and definitions. Three RSA models (target, distractor, incongruency) were fitted per region, yielding a set of 72 coding-strength estimates. These 72 estimates were used to predict Stroop interference effects within a cross-validated model selection procedure. **C**, Additional brain–behavior associations identified via cross-validated model selection. Left ventral visual incongruency coding (left panel) and left DLPFC distractor coding (right panel) were selected — in addition to right DLPFC target, right LPPC target, and left DMFC incongruency. See caption **A** for description of inset text, lines, and bands. **D**, Estimate of prediction accuracy of the selected model. Within a validation (held-out) set of 17 subjects (monozygotic co-twins of subjects within the analysis sample), observed Stroop effects were moderately correlated (red line) to predictions from the selected model. A permutation test (grey distribution) indicated the selected model significantly explained validation-set variance, suggesting the selected model results are relatively stable.

#### 2.3.11 Model Selection

To complement our hypothesis-driven brain–behavior analyses, we used a more data-driven model-selection approach. An expanded set of 24 superparcels (in addition to our six ROIs) was defined (Figure 3, B; see Figure 3-3 for list). Some superparcels were included as regions of interest, others were included as negative controls (i.e., regions that were not predicted to be important for explaining behavioral performance). Subject-level coefficients of the Stroop interference effect contrast were extracted from the behavior-only RT model and used as the response vector (in the model equation, the slope elements of **u***_s_*; also known as BLUPs or conditional modes). RSA was conducted on each superparcel (see *Estimation of Coding Strength β*), furnishing three *β* coefficients per superparcel. These 72 measures were fitted to the response vector via elastic net regression, implemented via *glmnet::glmnet()* in R (Friedman, Hastie, & Tibshirani, 2010). Parameter *α* was set to 0.5 (balancing lasso and ridge penalties). The parameter *λ* was tuned via 10-fold cross-validation (default) using the “1-SE rule”: the smallest *λ* greater than one SE of the minimum *λ* across folds was saved (Hastie, Tibshirani, & Friedman, 2009); this routine was repeated 1,000 times and the minimum mode (there were ties) of the saved *λ*s was selected. We selected the minimum mode because the maximum suppressed all variables from the model.

To assess validation-set accuracy, the selected model coefficients were applied to a design matrix from validation-set subjects, generating a predicted Stroop effect vector. The linear correlation was estimated between this predicted Stroop effect and observed Stroop effects (estimated as conditional modes via a hierarchical model separately fitted to validation-set data). The significance of this correlation was assessed by randomly permuting the training-set response vector, re-fitting the model, generating new predicted validation-set values, and re-estimating the predicted–observed correlation 10,000 times. The p-value was given by the proportion of resamples in which the null correlation was greater than the observed correlation.

#### 2.3.12 Exploratory Whole-Cortex RSA

The RSA-model fitting procedure, as outlined in *Estimation of Coding Strength β* was separately conducted on each MMP cortical parcel. Inferential statistics followed those suggested by Nili et al. (2014). One-sided signed-rank tests were conducted for significance testing (*>*0). P-values were FDR-corrected over all 360 parcels, separately for each task dimension (target, distractor, incongruency).

#### 2.3.13 Univariate Activation Analyses

We additionally conducted a standard “univariate activation” analysis on these data. Note that this was not meant to evaluate whether univariate activity was a plausible confounding variable in our analysis, but rather to provide some basis for comparing our data to most extant neuroimaging studies of Stroop. For a given ROI (or MMP parcel), beta coefficients from the first-level fMRI GLM were averaged over voxels by stimuli, then over stimuli by congruency. These mean values were then contrasted, analogous to the behavioral Stroop interference effect (incongruent − congruent). This statistic gives an estimate of the overall (across-voxel) difference in fMRI activity within a give brain region on incongruent versus congruent trials.

### 2.4 Follow-up Control Analyses

To establish the robustness of our results, we conducted several control analyses that examined a number of confounds and concerns: potential differences in signal-to-noise ratio across prefrontal ROIs, the effects of head motion, different RSA models for incongruency coding, the presence of bias imposed by the experimental design, within-run versus between-run RSA estimation, and the effects of downsampling to account for different trial numbers across runs. These are each described next.

#### 2.4.1 Comparison of Signal-to-Noise Ratios

To test for differences in SNR (signal-to-noise ratios) between DMFC and DLPFC, we estimated “noise ceilings” within each region and contrasted them across regions. Noise ceilings indicate the maximum observable group-level effect size (RSA model fit) given the level of between-subject variability in similarity structure (Nili et al., 2014). Lower (smaller) average noise ceilings indicate poorer SNR for group-level tests. We used the cross-validated “lower-bound” noise ceiling estimator of Nili et al. (2014), as this yields a lower-variance estimate, and therefore more powerful contrast across regions, than the non-cross-validated “upper-bound” (Hastie, Tibshirani, & Friedman, 2009, Section 7.10.1). For a given region, the lower-bound noise ceiling is defined for each subject *s* in 1*, …, N* as Cor(**y***_s_*, 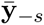): the linear correlation between a subject *s*’ observed similarity vector, **y***_s_*, and the group-mean vector excluding subject *s*, 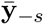. (See *Estimation of Coding Strength β* for definition of **y**.) As each of the *N* estimates are interdependent, we used a percentile bootstrap to contrast noise ceilings across regions. In each resample (of 10,000), noise ceilings were artanh transformed, contrasted across regions within-subject, then averaged across subjects. A two-sided p-value was provided by computing the proportion of resampled means greater than zero, *p*, then taking the minimum of 2*p* and 2(1 *−p*). Finally, we conducted “two one-sided tests” for equivalence (Lakens, 2017; Schuirmann, 1987) to affirm a null hypothesis of no difference between regions in noise ceiling. This consists of defining a threshold effect size, the minimum effect size of interest, and testing whether the observed difference in noise ceilings is significantly less extreme than the threshold. For an objective threshold, we used the smallest standardized effect size at which our bootstrap procedure was expected to retain 80% power, Cohen’s d = 0.35, determined via Monte Carlo simulation.

#### 2.4.2 RSA on Head Motion Estimates

As a negative control analysis for our exploratory whole-cortex RSA, we attempted to decode task variables (target, distractor, incongruency) via RSA from framewise estimates of head motion. The 6 motion regressors that were used in the fMRI GLM as nuisance covariates (corresponding to translation and rotation in 3 dimensions) were regressed upon the design matrix containing predicted BOLD timecourses of our 16 conditions of interest. The coefficient matrix resulting from this regression was then submitted to the RSA procedures described in *Estimation of Coding Strength β* and *Exploratory Whole-Cortex RSA*. To check whether more aggressive movement denoising within the fMRI GLM was warranted (i.e., in addition to the 6 nuisance regressors), we conducted this same movement-based RSA, however using 12 motion regressors (the 6 bases and their temporal derivatives). RSA model fits between the 6 and 12-basis motion-based RSAs were compared via paired-sample signed-rank test.

#### 2.4.3 Alternative RSA Incongruency Models

The RSA incongruency model parameterized the congruent–congruent correlations (i.e., *R_ij_* where *i* and *j* are both patterns from congruent trials, denoted here simply as *CC*) with a separate nuisance regressor. That is, these cells were effectively excluded, and the model instead computed the contrast *II − IC*. This exclusion was done because we have no specific hypotheses regarding how congruent trials should be encoded relative to one another. Other parameterizations are possible, however, including models that (a) incorporate *CC* correlations within the “baseline” or intercept term, by omitting the congruent nuisance regressor [i.e., *II −* ave(*IC* + *CC*)], or (b) that omit *IC* correlations from the contrast (i.e., *II − CC*). We note, however, that (b) is a suboptimal parameterization as it effectively excludes 40% (48/120) of observations per subject (i.e., all *IC* cells). To verify that our results were not dependent on the particular modeling approach we chose, we compared the observed RSA model fits to these alternative parameterizations. In brief, subject’s model fits were highly similar between the parameterizations (see **Results** sections *Group* and *Exploratory Whole-Cortex*; Figure 2-5). Thus, to help streamline the Results, we only conducted and reported analyses using the original RSA incongruency parameterization (in which the congruency model was a nuisance covariate).

#### 2.4.4 Design Bias

For the current study, a typical within-run form of RSA estimation was implemented, in which correlations were computed among activation patterns estimated within the same scanning run and first-level GLM. Within-run RSA has been criticized because it is susceptible to design biases that occur when trial orders are insufficiently randomized within the experiment (Cai, Schuck, Pillow, & Niv, 2019). A priori, this was not a strong concern in the current design, as trial orders were fully randomized both within and between participants. Nevertheless, we conducted several diagnostic and robustness analyses to validate that our results and conclusions were not impacted by this potential bias. First, we estimated the extent of the potential bias by conducting our RSA technique on data simulated under a “worst-case” scenario, that is, when SNR = 0 (details described within Figure 1-2) and across a wide range of autocorrelation strengths. At worst, the three models were weakly biased (within 0.02 to 0.05 of *α* = 0.05; Figure 1-2). This result indicated that while the bias was present, it was relatively minimal (cf., Cai, Schuck, Pillow, & Niv, 2019). Second, we validated that these simulated estimates were realistic, by using our actual fMRI data to estimate the false positive rate empirically. To do this, we conducted RSA on the first-level GLM coefficients from each subjects’ ventricles, as these voxels should contain no brain activity signal but similar noise characteristics as those of interest (a ventricle mask of 2431 voxels was obtained from AFNI servers: https://afni.nimh.nih.gov/pub/dist/tgz/suma_MNI_N27. tgz). By treating the group-level mean and SD of these RSA model fits as the parameters of a non-central null distribution, *X ∼ N* (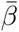*, SD*(*β*)), we computed the empirical false positive rate as *P* (*X* > *z*_*α*=0.05_), the proportion of this distribution greater than the customary one-tailed z-criterion. All rates were within 0.03 of *α* = 0.05 (target = 0.07, distractor = 0.04, incongruency = 0.08), confirming the bias was quite minimal.

#### 2.4.5 Between-Run RSA

As an alternative to within-run RSA, various between-run estimation approaches have been proposed which have been shown to be less sensitive to potential design biases. We opted not to use between-run RSA for our primary analyses, both because of the reduced effects of design bias established above, but also because between-run RSA is noted to be considerably more conservative that within-run RSA (Cai, Schuck, Pillow, & Niv, 2019). Moreover, several particulars of the present design are known to further hamper its power. Namely, between-run RSA makes incomplete use of the data, an issue which is exacerbated to the maximum extent possible in the present case, as our design has only the minimum number (two) of cross-validation folds (runs; Diedrichsen, Berlot, Mur, Schütt, & Kriegeskorte, 2020). Additionally, because the image acquisition sequence involved a reversal of phase-encoding direction across the two runs, this effectively effectively adds a strong non-linear component of noise if between-run RSA is used.

Nevertheless, to examine further the extent of design bias in our data, as well as the robustness of our results to the drop in power imposed by between-run RSA, we conducted a follow-up analysis of our primary results using between-run RSA approaches. We conducted two forms of between-run RSA: the first used the cross-correlation of patterns between scanning runs (“cross-run RSA”; see Alink, Walther, Krugliak, Bosch, & Kriegeskorte, 2015), and the second used a cross-validated form of the Euclidean distance (i.e., on the inner product of pattern contrasts between runs; “cross-validated RSA”; see Walther et al., 2016). We selected these two forms of RSA as they have complementary benefits. Cross-run correlation is most comparable to our original within-run correlation, as they are both linear correlations. However, using this method within our dataset also necessitated employing downsampling (as the number of trials per condition were not perfectly balanced at the run-level; see *Downsampling Analysis*), which increases the variance of resulting estimates due to discarding data. In contrast, cross-validated RSA is insensitive (in terms of expected value) to the issue of trial numbers per condition (Diedrichsen, Berlot, Mur, Schütt, & Kriegeskorte, 2020). Using this method therefore allowed us to conduct the RSA using all the data at once, without downsampling. But, cross-validated RSA tests a more constrained hypothesis than cross-run RSA. Whereas cross-run RSA can be sensitive to non-linear differences between conditions, cross-validated RSA tests linear discriminability between conditions. Thus, when a non-linear boundary separates conditions, cross-validated RSA will fail to detect an effect whereas cross-run RSA could succeed. A non-linear boundary would occur, for example, when one condition (e.g., incongruent stimuli) drives a reliable, common, response while the other conditions (e.g., congruent stimuli) either drive unreliable, or stable but heterogeneous, responses. Nevertheless, to make cross-validated RSA as comparable as possible to our primary RSA, we z-score standardized patterns prior to computation and omitted spatial prewhitening (Walther et al., 2016).

#### 2.4.6 Downsampling Analyses

Lastly, we checked whether the primary results were robust to the prewhitening method of data preprocessing, which was introduced to handle the imbalance of trials across runs (see *Method* section *Estimation of coding strength β*, step 4). In this analysis, we instead handled this issue by performing RSA after equating the number of trials per run*condition, by iteratively downsampling conditions with random subsets of trials. Specifically, we first fitted GLMs on fMRI timeseries, separately for each scanning run, that contained a single regressor per trial (LS-A method of Mumford, Turner, Ashby, & Poldrack, 2012). The minimum number of times we presented each unique Stroop stimulus in a single run was 3; this was the number to which we downsampled all conditions with *>*3 occurrences. For these conditions, we randomly sampled three trials, and averaged GLM coefficients voxel-wise over these trials. (For 3-trial conditions, we simply averaged all trials that were present.) This formed 16 separate condition-level coefficient vectors (activity patterns) per run. We then averaged these coefficient vectors across run, then estimated condition*×*condition correlation matrices from these patterns (averaging across runs was omitted from the downsampled cross-run RSA; see preceding section). We repeated this resampling, averaging, and correlation process 1,000 times, and averaged the resulting correlation matrices across iterations (after an artanh transform). These correlation matrices were then submitted to the same RSA as outlined in *Estimation of coding strength β*, with the omission of the prewhitening step (4).

## 3 Results

Influential theories of cognitive control have proposed specific dissociations in the type of task information encoded by human medial and lateral frontoparietal cortex (Figure 1, A–B). But, previous studies have largely approached this question indirectly, by employing tasks designed to recruit these regions differentially in time, then testing for temporal dissociations in regional-mean levels of fMRI activity. Here, we used a more direct approach, by using the similarity structure of neural activity patterns evoked within these regions to estimate their informational content. In particular, through RSA, we compared neural coding of three distinct types of Stroop-task information — target, distractor, and incongruency (Figure 1, D–E) — within each region of interest (ROI) simultaneously, while Stroop interference was being experienced and resolved.

We describe three sets of analyses. First, we examined group-level effects, to test for neuroanatomical dissociations in encoding of task information (Figure 1, B, center). Second, we examined individual-level effects (i.e., individual differences), to test for dissociations in brain–behavior relationships (Figure 1, B, right). These two analyses were ROI-based and primarily focused on dorsomedial frontal and lateral frontoparietal regions (Figure 1, D), but also included sensorimotor ROIs for comparison purposes. The last set of analyses was conducted in whole-cortex exploratory fashion, to provide a more comprehensive picture of the anatomical profile of each task dimension.

### 3.1 Group

#### 3.1.1 Dorsomedial frontal and dorsolateral prefrontal cortex exhibit distinct coding profiles

Primary group-level results are summarized in Figure 2. Statistical estimates corresponding to results outlined within this section are contained in Table 2.

**Table 2:** Group-level results from RSA in frontoparietal regions of interest. Group-mean RSA model fits and contrasts between RSA models (target, distractor, incongruency) and between ROIs (DMFC, DLPFC, LPPC in each hemisphere). Each row displays the statistical estimates from a given RSA model and ROI, or from a contrast between RSA models or ROIs. Contrasts were conducted either between RSA models (within ROI) or between ROIs (within RSA model). For example, target − distr. | DLPFC (R) indicates the contrast between target and distractor RSA model fits (‘coding strengths’) within right DLPFC. 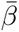 indicates the mean RSA model fit over subjects. Unless noted, contrasts were averaged across hemisphere. For contrasts performed separately in each hemisphere, p-values were FDR-adjusted for two comparisons. Products of bracketed terms indicate interactions between RSA model and ROI. Dark grey rows display contrasts with *p* < 0.05, light grey rows display contrasts with 0.05 < *p* < 0.1 Incon., incongruency; distr., distractor.

| contrast | $\bar{\beta}$ | $\sigma$ | t | p |
| --- | --- | --- | --- | --- |
| DMFC (L) incon. | 0.11 | 0.03 | 3.65 | 0.00026 |
| DMFC (R) incon. | 0.08 | 0.03 | 2.82 | 0.00485 |
| DMFC (L) target | 0.04 | 0.01 | 2.61 | 0.00918 |
| DMFC (R) target | 0.05 | 0.01 | 3.94 | 0.00008 |
| DMFC (L) distr. | 0.01 | 0.01 | 0.78 | 0.43667 |
| DMFC (R) distr. | 0.00 | 0.01 | 0.19 | 0.84907 |
| DLPFC (L) incon. | 0.04 | 0.02 | 2.04 | 0.04155 |
| DLPFC (R) incon. | 0.05 | 0.02 | 2.39 | 0.01682 |
| DLPFC (L) target | 0.03 | 0.01 | 2.56 | 0.01062 |
| DLPFC (R) target | 0.04 | 0.01 | 3.18 | 0.00147 |
| DLPFC (L) distr. | -0.01 | 0.01 | -0.98 | 0.32561 |
| DLPFC (R) distr. | 0.00 | 0.01 | 0.31 | 0.75797 |
| LPPC (L) incon. | 0.07 | 0.02 | 3.07 | 0.00211 |
| LPPC (R) incon. | 0.06 | 0.02 | 2.64 | 0.00818 |
| LPPC (L) target | 0.03 | 0.01 | 2.74 | 0.00617 |
| LPPC (R) target | 0.05 | 0.01 | 4.38 | 0.00001 |
| LPPC (L) distr. | 0.00 | 0.01 | 0.28 | 0.77722 |
| LPPC (R) distr. | 0.02 | 0.01 | 1.86 | 0.06359 |
| incon. – target \| DMFC (L) | 0.07 | 0.03 | 2.26 | 0.04737 |
| incon. – target \| DMFC (R) | 0.03 | 0.03 | 0.91 | 0.36047 |
| incon. – distr. \| DMFC (L) | 0.10 | 0.03 | 3.22 | 0.00256 |
| incon. – distr. \| DMFC (R) | 0.08 | 0.03 | 2.64 | 0.00826 |
| target – distr. \| DMFC (L) | 0.03 | 0.02 | 1.63 | 0.10230 |
| target – distr. \| DMFC (R) | 0.05 | 0.02 | 3.18 | 0.00297 |
| incon. – target \| DLPFC (L) | 0.01 | 0.02 | 0.46 | 0.65809 |
| incon. – target \| DLPFC (R) | 0.01 | 0.02 | 0.44 | 0.65809 |
| incon. – distr. \| DLPFC (L) | 0.06 | 0.02 | 2.41 | 0.03212 |
| incon. – distr. \| DLPFC (R) | 0.05 | 0.02 | 2.09 | 0.03671 |
| target – distr. \| DLPFC (L) | 0.04 | 0.02 | 2.76 | 0.01145 |
| target – distr. \| DLPFC (R) | 0.04 | 0.02 | 2.33 | 0.01989 |
| incon. – target \| LPPC (L) | 0.04 | 0.03 | 1.62 | 0.21226 |
| incon. – target \| LPPC (R) | 0.01 | 0.03 | 0.43 | 0.66707 |
| incon. – distr. \| LPPC (L) | 0.07 | 0.03 | 2.75 | 0.01202 |
| incon. – distr. \| LPPC (R) | 0.04 | 0.03 | 1.63 | 0.10282 |
| target – distr. \| LPPC (L) | 0.03 | 0.02 | 1.87 | 0.06185 |
| target – distr. \| LPPC (R) | 0.03 | 0.02 | 1.96 | 0.06185 |
| DMFC (L) – DLPFC \| incon. | 0.06 | 0.02 | 2.38 | 0.01737 |
| DMFC (L) – DLPFC \| target | 0.00 | 0.01 | -0.09 | 0.92467 |
| DMFC (L) – LPPC \| incon. | 0.04 | 0.03 | 1.47 | 0.14112 |
| DMFC (L) – LPPC \| target | -0.01 | 0.01 | -0.56 | 0.57868 |
| [incon. – target] · [DMFC (L) – DLPFC] | 0.06 | 0.03 | 2.20 | 0.02797 |
| [incon. – target] · [LPPC (L) – DLPFC] | 0.03 | 0.03 | 1.06 | 0.29066 |

The DMFC has been strongly associated with the coding of incongruency information in response conflict tasks such as the Stroop. RSA approaches were used to directly test the specificity of that hypothesis. This region was indeed found to encode the incongruency dimension of the Stroop task (left: 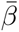 = 0.11*, p* = 0.00, right: 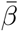 = 0.08*, p* = 0.00). Further, within left DMFC, a preference for this dimension was observed: incongruency information was encoded more strongly than either target or distractor information (vs. target: Δ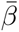 = 0.07*, p* = 0.05; vs. distractor: Δ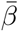 = 0.10*, p* = 0.00; p-values adjusted across hemisphere; Figure 2, A). This DMFC preference was prominent enough that it could be seen in its overall similarity structure within a low-dimensional embedding (Figure 2, B). Further, incongruency coding was neuroanatomically dissociated, as this coding scheme was reflected more strongly in left DMFC than in DLPFC activity patterns (Δ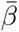 = 0.06*, p* = 0.02; Table 2).

We next focused on lateral frontoparietal regions and the coding of target information. DLPFC indeed encoded target information (left: 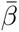 = 0.03*, p* = 0.01 right: 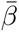 = 0.04*, p* = 0.00), and more strongly than distractor information (Δ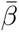 = 0.04*, p* = 0.00). We did not find a preference, however, for target over incongruency information in DLPFC (Δ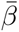 = −0.01*, p* = 0.62), nor did we observe significantly stronger target coding than in DMFC (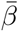 = 0.00*, p* = 0.92). A qualitatively similar pattern of results was observed in lateral posterior parietal cortex: significant target and incongruency coding, significantly enhanced target versus distractor coding, but no observed preference among target or incongruency nor detectable difference from DMFC target coding (Table 2).

Thus, at the group-level, we observed a single rather than double dissociation between medial and lateral frontoparietal cortex, in the form of enhanced sensitivity to incongruency relative to target coding in left DMFC. In all ROIs, however, these two sources of task information were more strongly encoded than distractor information.

#### 3.1.2 Sensitivity and control analyses

We next tested a series of hypotheses to scrutinize and extend our results.

First, we conducted a positive control analysis to bolster confidence in the statistical power of RSA methods within the present design. In particular, we sought to determine whether our methods could detect dissociations in task coding that are strongly expected to exist. For this, we focused on primary somatomotor and visual cortical ROIs, the responses of which can be assumed to reflect, relatively selectively, response-related (i.e., motoric) and visual form-related coding. As the distractor (word) defines the visual form of the Stroop stimulus, coding of form-related features should be captured by our distractor model. In parallel, as our analysis included only correct-response trials (i.e., in which the target response was spoken), coding of motoric features should be captured by our target model. Consistent with this logic, within early visual cortex, evidence of preferential distractor coding was observed (Figure 2, C; distractor: 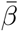 = 0.04, *p* < 0.001, target: 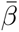 = 0.02*, p* = 0.17, distractor vs. target: Δ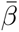 = 0.03*, p* = 0.05), whereas, within primary ventral somatomotor cortices (encompassing the “mouth” homunculi), a relatively selective pattern of target coding was found (Figure 2, C; distractor: 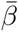 = 0.00*, p* = 0.9, target: 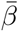 = 0.13, *p* < 0.001, distractor vs. target: Δ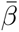 = −0.13, *p* < 0.001). Further, coding of these dimensions was strong enough to predominate the overall structure of patterns within a low-dimensional embedding (Figure 2, B). Thus, in primary visual and somatomotor regions, a relatively clear-cut group-level double-dissociation emerged. This suggests that our models were adequately powered, at least in primary sensorimotor cortices, to detect functional distinctions, and that the failure to observe distractor coding in DMFC and DLPFC was not due to a general deficiency in our distractor coding model.

Second, we conducted sensitivity analyses to assess the robustness of our results to the particular ROI definitions used. In one analysis, we tested more expansive ROI definitions, by using alternatively-defined superparcels (Figure 1-1). These definitions included additional, more rostral PFC parcels (1 in DMFC, 3 in DLPFC), which begin to encroach into ventromedial PFC and frontopolar cortex (e.g., the rostral DMFC parcel was assigned to the Default-Mode network within the Cole-Anticevic divisions). Nevertheless, the previously observed dissociations were robust to these more liberal definitions (Figure 2-1). In the other analysis, we examined whether the overall superparcel coding profiles were representative of individual parcels. While DMFC and DLPFC results generally reflected that of constituent parcels (Figure 2-2, A, B), interestingly, there was substantial heterogeneity within left LPPC (Figure 2-2, C). Similar to left DMFC, a collection of left LPPC regions spanning the length of intraparietal sulcus strongly encoded the incongruency dimension (i.e., IP1, IP2, IPS1, AIP, LIP, MIP).

Third, we examined whether the lack of observed discrimination between target and incongruency coding dimensions within DLPFC could be explained by increased error variance potentially present in fMRI activity patterns within this region. Prior work has suggested that PFC regions might be particularly susceptible to this confound (Bhandari, Gagne, & Badre, 2018). It is possible that we might have observed a dissociation in left DMFC but not DLPFC due to differential levels of statistical power across the two regions. We therefore derived an SNR (signal-to-noise ratio) analysis to determine whether this was a viable explanation (see Method). A paired-sample bootstrap test did not indicate a systematic difference between DLPFC versus left DMFC group-level SNR (Δ*z* = −0.01*, p* = 0.60). However, the SNRs in these regions were also not confirmably similar, as indicated by an equivalence test (which provides a confirmatory test of the null hypothesis; *p* = 0.10). Therefore, while we cannot rule out the possibility that the single dissociation observed between left DMFC and DLPFC was driven by better group-level SNR in DMFC, any potential SNR differences between the two regions were not substantial enough for our methods to detect.

Fourth, to provide a basis for comparison to most extant neuroimaging research of the Stroop task, we conducted a “univariate activation” analysis, examining whether these brain regions were generally more active during incongruent versus congruent conditions. No regions were found to respond more strongly overall to incongruent versus congruent conditions (Figure 2-3, A), although the mean contrast in DMFC (L) was positive (i.e., incongruent*>*congruent). This null result was not surprising, however, due to the high frequency of incongruent trials within the experiment — which is known to reduce both the behavioral and neural univariate Stroop effect (Carter et al., 2000; De Pisapia & Braver, 2006; Logan & Zbrodoff, 1979). While this null result demonstrates the utility of using RSA in this case, it should not, however, be seen as direct evidence for the increased sensitivity of RSA versus univariate methods, as the univariate and RSA-based tests as implemented here are subject to different constraints and are thus incomparable (Allefeld, Görgen, & Haynes, 2016). Finally, the magnitude of the univariate Stroop effects were only weakly correlated incongruency coding model fits (Figure 2-3, B), suggesting these measures were non-redundant.

Finally, we tested whether our pattern of results were robust to alternative RSA techniques, including a downsampling technique to equate trial counts across runs, two “between-run” RSA methods (cross-run RSA and cross-validated RSA; see Method), and alternative parameterizations of the incongruency coding model (see Method). Findings were robust to downsampling and to between-run RSA (Figure 2-4), and were highly similar across different parameterizations (Figure 2-5). Interestingly, however, the detection of incongruency coding in DMFC depended on whether a linear or non-linear RSA method was used. When using an RSA method that tests linear discriminability between conditions (cross-validated RSA), the incongruency coding effect was abolished in DMFC; whereas when using a comparable method that is sensitive to non-linear pattern differences (cross-run RSA), the effect remained quite strong. This pattern of results suggests that incongruency information was encoded non-linearly within DMFC activation patterns. Indeed, this can be seen within the 2-dimensional embedding (Figure 2), as a radial, rather than linear, separation of incongruent (central) and congruent (peripheral) stimuli.

### 3.2 Individual

Primary individual-level results are summarized in Figure 3. Statistical estimates corresponding to results outlined within this section are contained in Table 3 (see Figure 3-1 for scatter plots of all associations).

**Table 3:** Parameter estimates from hierarchical brain–behavior models that explained individual differences in behavioral performance (RT) with variability in the strength of neural coding (*β*, see Method). Separate models were fit per region and RSA model (target, incongruency) combination. The displayed estimates correspond to the interaction **stroop** · *β*, which indicates how subjects’ coding strengths relate to the magnitude of their Stroop interference effects. Also displayed are contrasts on **stro**op · *β* estimates, between RSA models (within ROI) or between ROIs (within RSA model). For example, target − incon. | DLPFC (R) indicates, for right DLPFC, whether target and incongruency coding were differentially related to the size of the Stroop interference effect. Dark grey rows display contrasts with *p* < 0.05, light grey rows display contrasts with 0.05 < *p* < 0.1

| contrast | b | $\sigma$ | t | p |
| --- | --- | --- | --- | --- |
| DMFC (L) incon. | 47.66 | 25.97 | 1.84 | 0.06647 |
| DMFC (R) incon. | -0.10 | 33.15 | 0.00 | 0.99762 |
| DMFC (L) target | 48.40 | 90.63 | 0.53 | 0.59330 |
| DMFC (R) target | -24.04 | 63.75 | -0.38 | 0.70613 |
| DLPFC (L) incon. | 14.08 | 41.17 | 0.34 | 0.73227 |
| DLPFC (R) incon. | 38.42 | 33.66 | 1.14 | 0.25367 |
| DLPFC (L) target | -105.93 | 78.34 | -1.35 | 0.17637 |
| DLPFC (R) target | -239.13 | 71.68 | -3.34 | 0.00085 |
| LPPC (L) incon. | -48.95 | 38.13 | -1.28 | 0.19921 |
| LPPC (R) incon. | -32.40 | 32.22 | -1.01 | 0.31467 |
| LPPC (L) target | -1.90 | 93.65 | -0.02 | 0.98385 |
| LPPC (R) target | -235.34 | 71.23 | -3.30 | 0.00096 |
| incon. – target \| DMFC (L) | 20.92 | 97.23 | 0.22 | 0.82962 |
| incon. – target \| DLPFC (R) | 299.21 | 80.62 | 3.71 | 0.00021 |
| incon. – target \| LPPC (R) | 204.56 | 79.94 | 2.56 | 0.01050 |
| DMFC (L) – DLPFC (R) \| incon. | 22.85 | 52.92 | 0.43 | 0.66587 |
| DMFC (L) – DLPFC (R) \| target | 500.63 | 139.14 | 3.60 | 0.00032 |
| DMFC (L) – LPPC (R) \| incon. | 134.19 | 49.85 | 2.69 | 0.00710 |
| DMFC (L) – LPPC (R) \| target | 468.99 | 137.46 | 3.41 | 0.00065 |

#### 3.2.1 Better-performing subjects have stronger lateral frontoparietal target coding

The fidelity of target-related information in lateral frontoparietal cortex — DLPFC, in particular — is thought to be closely linked to the efficiency with which an individual resolves response conflict in tasks such as Stroop. By using subject-level target-coding estimates (*β*_target_) to model behavioral performance (RT), we tested this fundamental prediction relatively directly. Indeed, subjects with stronger target coding in both right DLPFC and right LPPC resolved Stroop interference effects more quickly (DLPFC: *b* = −239.13*, p* = 0.00*, r* = −0.44 [−0.66*, −*0.15]*, ρ* = −0.39 [−0.63*, −*0.08]; LPPC: *b* = −235.34*, p* = 0.00*, r* = −0.44 [−0.63*, −*0.17]*, ρ* = −0.35 [−0.59*, −*0.06]; p-values corrected across hemisphere; linear correlation, *r*; rank correlation, *ρ*; bootstrapped 95% confidence interval, [*lower, upper*]; Figure 3, A; Table 3). Subjects’ target coding estimates were moderately correlated (*r* = 0.53) between these two lateral frontoparietal regions, as expected based on their strong neuroanatomical and functional connectivity (e.g., Buschman & Miller, 2007; Petrides & Pandya, 1999). In neither of these lateral regions was incongruency coding significantly related to the Stroop interference effect (DLPFC: *b* = 38.42*, p* = 0.25*, r* = −0.17 [−0.14, 0.45]*, ρ* = −0.10 [−0.19, 0.37]; LPPC *b* = −32.40*, p* = 0.31*, r* = −0.15 [−0.40, 0.13]*, ρ* = −0.18 [−0.45, 0.12]; p-values uncorrected), and within right DLPFC, target coding was a considerably stronger explanatory variable than incongruency coding (Δ*b* = −299.21*, p* = 0.00; Table 3).

Conversely, incongruency-related responses of DMFC are thought to be positively associated with maladaptive policies of response selection. In line with this notion, subjects with stronger incongruency coding in left DMFC tended to exhibit greater Stroop interference, although this interaction was non-significant (*b* = 47.66*, p* = 0.13*, r* = −0.26 [−0.01, 0.49]*, ρ* = −0.17 [−0.12, 0.44]; p-value corrected across hemisphere; Figure 3, A). Similarly, incongruency coding was numerically more strongly associated with Stroop interference effects than target coding in this region, though this statistical difference was also non-significant (Δ*b* = −20.92*, p* = 0.83; Table 3). While weak, this DMFC–Stroop association notably emerged within the same hemisphere that displayed a group-level preference for incongruency information (Figure 2, B). Importantly, the sign of the correlations we detected within our ROIs were robust to alternative RSA techniques (Figure 3-2; note that general attentuation in effect size is expected with higher variance techniques; see *Downsampling Analysis* and *Between-Run RSA* in Method).

Considering the collective pattern of results, we conclude here that our findings support the hypothesis that target coding in (right) DLPFC and in LPPC reflected a common process of implementing control.

#### 3.2.2 Model selection affirms a lateral–medial dissociation and identifies unexpected relationships

Because the preceding hypothesis-driven analysis exclusively focused on a limited set of regions, important brain–behavior relationships may have been missed. A more accurate model may even omit target and incongruency coding from DLPFC, DMFC and LPPC altogether. Consequently, we conducted a more comprehensive test to identify regions and task dimensions that could better account for Stroop performance variability across individuals.

A data-driven model selection analysis was conducted to address this question (see Method). We defined an expanded set of 24 cortical regions (superparcels), including the six defined and used earlier, which covered various areas that may be important for performing the Stroop task (Figure 3, B; Figure 3-3). Conducting RSA on each superparcel furnished three coding estimates (one per coding model) per superparcel. These 72 estimates were then used as features in a cross-validated model selection procedure.

Strikingly, the selected model contained all three hypothesized measures: (right) DLPFC and LPPC target coding, and (left) DMFC incongruency coding. In addition, two unexpected measures were identified, both with negative slopes: left DLPFC distractor coding (*r* = −0.43 [−0.66*, −*0.14]*, ρ* = −0.33 [−0.60*, −*0.04]) and left ventral visual incongruency coding (*r* = −0.35 [−0.52*, −*0.13]*ρ* = −0.43 [−0.62*, −*0.20]; bivariate correlations shown in Figure 3, C). We tested the predictive accuracy of the selected model by using a held-out validation set of 17 subjects (co-twins of the primary analysis set). The selected model explained significant variance in Stroop interference effects within the validation set (Figure 3, D). While not fully independent, these validation set data were from a true hold-out and obtained from distinct individuals. This result therefore bolsters claims regarding the stability of the selected model.

Nevertheless, to provide a cursory test of a truly independent validation set (i.e., with no familial dependency to the training set), we excluded all subjects in the training set who were co-twins of with those in the validation set, and reconducted this model selection procedure. This amounted to discarding 16/49 (33%) of training-set observations. The selected model contained only one variable, which was not in our ROIs (but in early visual cortex), and was unable to predict held-out Stroop effects (*r* = −0.12). This is perhaps unsurprising, however, given the substantial reduction in the size of the training dataset for an already high-dimensional model. To reduce the dimensionality, we reconducted this analysis, focusing now instead only on ROIs and coding schemes of interest — target coding in DLPFC and LPPC, and incongruency coding in DMFC (within each hemisphere) — and additionally ensured that all variables were used in prediction (via ridge regression). This model was better able to predict the held-out Stroop effect (*r* = 0.20), in particular, relative to a comparable model that contained theoretically “mismatched” ROI*×*coding-scheme combinations (incongruency coding in DLPFC and LPPC, target coding in DMFC; *r* = −0.17, bootstrapped *p* = 0.088).

Results from these model selection analyses affirm a functional dissociation across the medial– lateral axis of frontoparietal cortex, and further demonstrate that Stroop-task representations within DLPFC and DMFC hold relatively privileged relationships with behavior.

### 3.3 Exploratory Whole-Cortex RSA

In a final, exploratory analysis, we estimated RSA models separately for each MMP parcel, to determine more comprehensively how target, distractor, and incongruency coding are distributed across cortex. These three task dimensions were encoded across cortex according to different neuroanatomical profiles. Target coding was widespread (observed in 207/360 parcels), covering substantial portions of the frontal and temporal lobes, including many perisylvian regions (Figure 4, top; Figure 4-1). Notably, the strongest target coding was observed within regions that receive strong sensory and (or) motor-related input (Figure 4-1). Contrastingly, incongruency coding was detected predominantly within prefrontal and intraparietal sulcal parcels — including DMFC, but also left LPPC, bilateral superior frontal gyrus, and left lateral frontopolar cortex (rostral DLPFC) — but additionally within left retrosplenial and right lateral occipital cortex (Figure 4, middle; Figure 4-3). Aside from this occipital area, these incongruency-coding parcels notably belonged to control networks (frontoparietal, cinguloopercular, and dorsal attention; Figure 4-3). In a third, distinct profile, distractor coding was only observed within early visual cortex (left V1 and V2; Figure 4, bottom; Figure 4-2).

**Figure 4:**
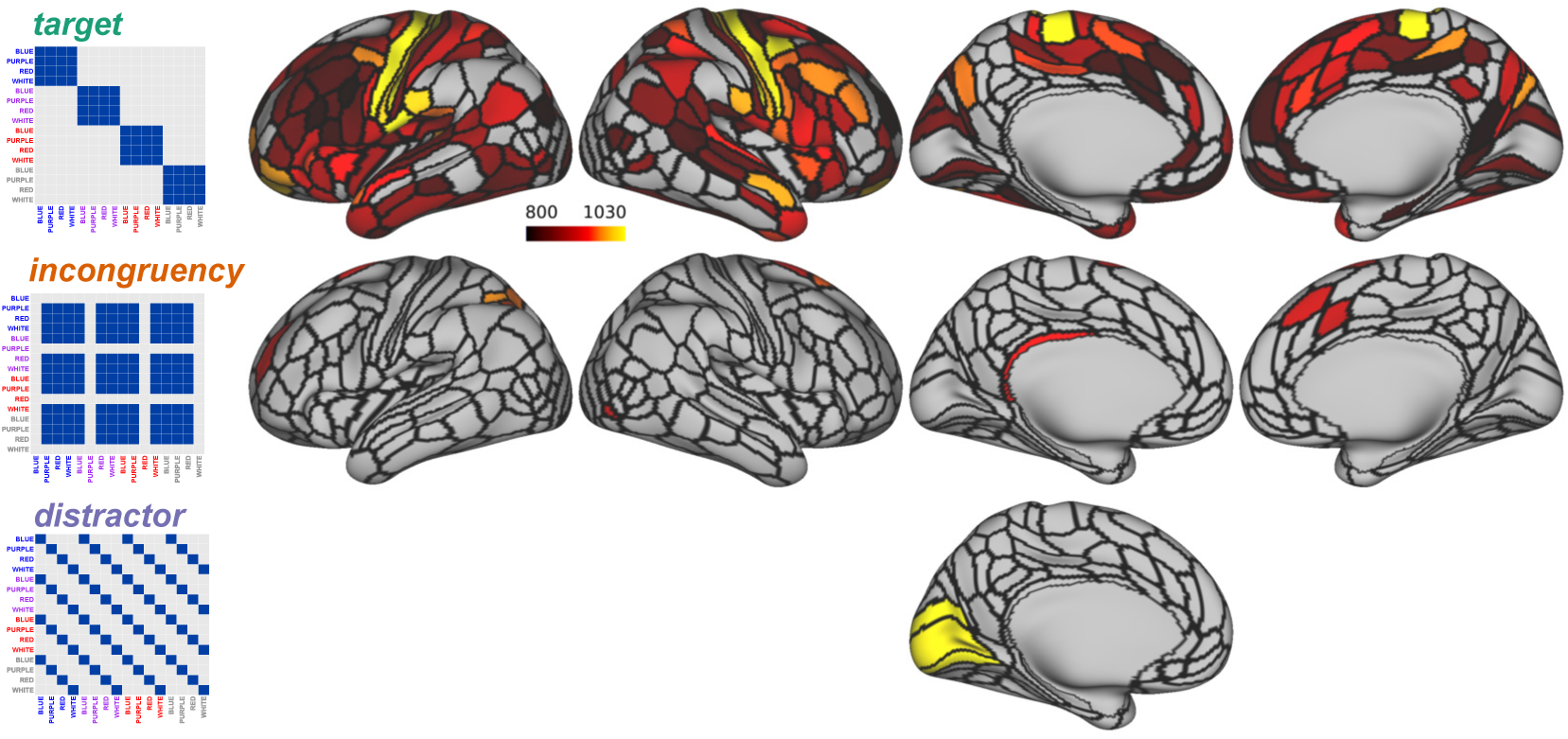
Cortical distributions of target (top), incongruency (middle) and distractor (bottom) coding, identified via an exploratory, whole-cortex analysis. Color bar indicates test statistic from one-sided sign-ranked test. For all statistical estimates, see Figures 4-1–4-3. Sensitivity and robustness analyses suggested that the core results were robust to various analysis decisions and implementations (handling of head motion, see text in Results; parameterization of incongruency coding, Figure 2-5, B; RSA estimation method, Figure 4-4).

As a negative control analysis, we tested whether we could decode these three task variables (target, distractor, incongruency) from framewise head motion estimates, using the same RSA procedures as above. No task variable was significantly encoded within patterns of head movements (*b*s < 0.01, *p*s *>* 0.36). Using a larger basis set of motion estimates (12 vs 6) did not yield significant decoding (*b*s < 0.01, *p*s *>* 0.17), nor any increased sensitivity to task variables (*b*s < 0.00, *p*s *>* 0.17), suggesting that our motion removal procedures (scrubbing and 6 motion regressors) were adequate.

Finally, we repeated this exploratory analysis using alternative RSA methods involving downsampling and between-run RSA (*Downsampling Analysis*, and *Between-Run RSA* sections of Methods; Figure 4-4). Across these analyses, the core results were quite robust: we found incongruency coding in parcels within DMFC (though this depended on the use of non-linear RSA methods, as in Figure 2-4 and Figure 3-2), target coding in mid-DLPFC, and distractor coding in visual cortex. Our findings were therefore not specific to a particular estimation method.

Collectively, these exploratory results confirm and extend our prior findings. (1) As with the reported brain–behavior associations, target coding was emphasized relative to coding of other task dimensions. (2) Yet despite this emphasis, important and expected dissociations in anatomical profiles were identified across our three coding models, further suggesting that these models were successful in measuring coding of distinct task dimensions (Figure 1, A).

## 4 Discussion

We analyzed the similarity structure, or representational geometry (Kriegeskorte & Kievit, 2013), of frontoparietal activity patterns associated with cognitive control, during performance of the classic color-word Stroop task. In left DMFC, incongruency coding predominated. While DLPFC and LPPC encoded both target and incongruency-related information, distractor coding was not detected in these regions, but was instead identified in early visual cortex. Further, these neural coding estimates were important and specific indicators of individual differences in magnitude of the behavioral Stroop interference effect. Individuals with stronger target coding in right DLPFC and right LPPC, but weaker incongruency coding in left DMFC, exhibited enhanced cognitive control, in terms of a reduced Stroop effect. Further, in a more comprehensive predictive model that included coding measures from a wide set of cortical regions, coding measures specifically from lateral frontoparietal and dorsomedial frontal regions were privileged in their link to behavior.

On one level, this study is a specific extension of research that has drawn dissociations between control-related functions of DLPFC and DMFC (Floden, Vallesi, & Stuss, 2010; MacDonald, Cohen, Stenger, & Carter, 2000). Most prominently, MacDonald et al. (2000) used a modified Stroop-task design, in which task rules (delivered via pre-cues) randomly alternated between color naming and word reading across trials, to demonstrate that DLPFC and DMFC encoded different types of information during cognitive control engagement. In particular, DLPFC was selectively recruited following cues for the more demanding color-naming task, whereas DMFC was instead driven by incongruent color-naming trials. This pattern of recruitment suggested that DLFPC encodes task-set and rule-related information in a preparatory manner, whereas DMFC encodes incongruency and conflict-related information in a stimulus-evoked manner. In the current study, we leveraged the high spatial dimensionality of fMRI to test whether this functional dissociation can be observed *within* a common time window of response selection, and further with a more traditional Stroop-task design, which does not involve task cues or switches. Our findings reinforce the conclusions of these relatively low-powered studies (N = 12 in Flooden et al., 2010; N = 9 in MacDonald et al., 2000) and indicate that the dissociations were not dependent upon the use of cued task-switching designs. Synthesizing these prior findings with those of the present study hints at a continuity in the putative role of DLPFC during target selection. Rather than exclusively contributing to preparation, DLPFC coding may evolve from proactively representing abstract rule or set-related information, towards more concrete targets and behavioral choices as relevant stimulus information becomes available in the environment. This view accords with work in monkey electrophysiology (Mante, Sussillo, Shenoy, & Newsome, 2013; Rigotti et al., 2013; Stokes et al., 2013), yet further work is needed to determine whether similar dynamics occur within human DLPFC and how such dynamics may reflect or interact with specific processes of cognitive control in Stroop-like tasks.

More broadly, the results of this study highlight the utility of RSA and the general representational geometric framework for investigating cognitive control. Previous work has used MVPA decoding in the Stroop task to study the impact of control demand on posterior representations (e.g., Banich et al., 2019). Here, we used the RSA framework to explicitly model and decompose control-related frontoparietal representations. In fact, a major motivation of the current study was to assess how well RSA measures of frontoparietal coding map to theorized mechanisms of control. For this purpose, the medial–lateral functional dissociation in frontoparietal cortex was a useful test-bed, as it features in several theoretical accounts (Botvinick, Braver, Barch, Carter, & Cohen, 2001; Miller & Cohen, 2001; Ridderinkhof, Ullsperger, Crone, & Nieuwenhuis, 2004; Shenhav, Botvinick, & Cohen, 2013). Our results were generally in line with these accounts, joining with a growing body of research in suggesting that RSA provides a convenient yet powerful framework from which neural measures can be used to test cognitive control theory (see Freund, Etzel, & Braver, 2021, for a review).

Nevertheless, the current work represents only an initial step in using the RSA framework to investigate cognitive control within Stroop-like tasks. As such, our study raises a number of unaddressed questions. But promisingly, there are ample opportunities for improving and extending the RSA framework highlighted here. For instance, a key limitation of the current study was the finding of widespread coding of the target dimension, suggesting a lack of specificity in the target RSA model. This is perhaps not surprising, however, as the model would capture not only coding of attentional-template and choice-related information, but also hue and response-related information. We mitigated this issue by demonstrating that target coding was selectively related to behavior within DLPFC and LPPC. Yet, this limitation could be addressed more powerfully by experimental design. Adding specific factorial manipulations, such as a task rule manipulation (*a la* MacDonald, et al. 2000; Hall-McMaster, Muhle-Karbe, Myers, & Stokes, 2019), or a response modality manipulation (*a la* Minxha, Adolphs, Fusi, Mamelak, & Rutishauser, 2020; see also Barch et al., 2001), would enable a richer, more precise set of cognitive control-relevant coding variables to be estimated (Figure 5).

**Figure 5:**
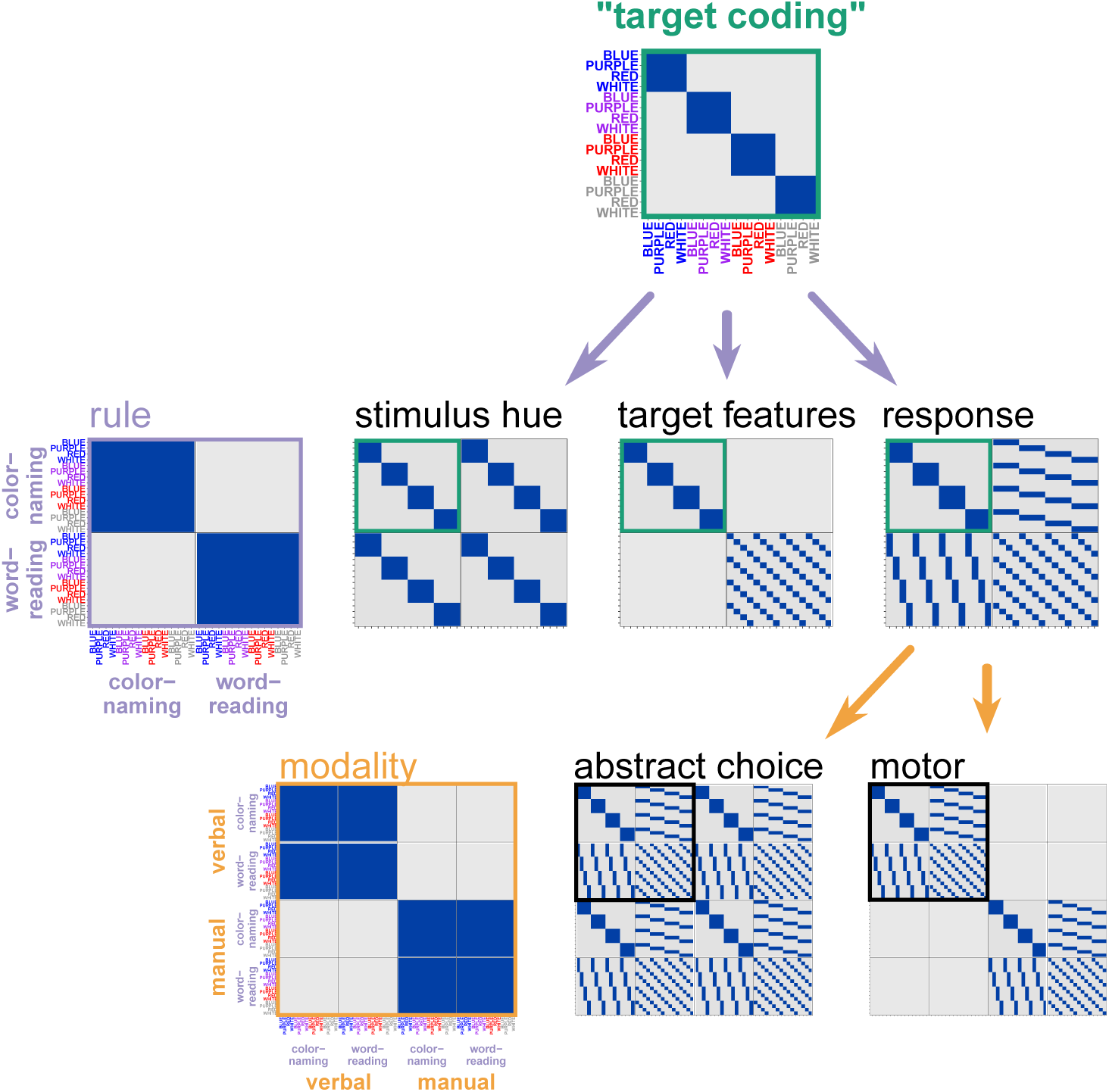
Expanded experimental designs. The current target coding model (top, green text; see also Figure 1) conflates distinct coding schemes, including those associated with relatively ‘early’ vision, relatively ‘late’ motoric commands, or more “central” rule-dependent schemes. By adding a factorial manipulation of task rule (i.e., participants respond to the present set of stimuli, but under both color-naming and word-reading conditions; MacDonald, et al. 2000; light purple arrows and text), the target coding model can be expanded into three more precise models (middle row of matrices), which are identical to our “target coding” model within the top-left quadrant (green square; naming–naming), but are now distinguished among the other quadrants (reading–reading, reading–naming). “Stimulus hue” and “response” models would identify coding that depends upon features of either the stimulus or response, independent of the task rules. Conversely, “target features” would identify a more flexible “attentional template” coding scheme, which depends critically on the current task goals (note that, the lower-right quadrant resembles our “distractor” coding model in Figure 1, as word features become the target in the word-reading condition). Importantly, too, abstract coding of task rule would also now be distinguishable (“rule” matrix on left). For an elegant example of a similar design within cued-task switching, see Hall-McMaster, Muhle-Karbe, Myers, and Stokes (2019). The “response” model could be further elaborated by incorporating another manipulation of response modality (verbal, manual; orange arrows). Now, coding of abstract choice options, independent of effector, could be separated from effector-dependent motoric coding. For an elegant example of a similar manipulation, see Minxha, Adolphs, Fusi, Mamelak, and Rutishauser (2020).

Future work could also address some of the complexities revealed by our data that were not entirely accounted for by the theoretical frameworks we used. For one, although predicted coding profiles emerged in some frontoparietal regions of interest, all regions encoded incongruency and target information. This incongruency-coding finding is consistent with prior univariate fMRI research (e.g., Nee, Wager, & Jonides, 2007; Niendam et al., 2012), and a more recent finding that the response of single neurons in human dACC and DLPFC are robustly modulated by conflict (Smith et al., 2019). With respect to target coding, however, one speculative interpretation is that, during the relatively late phase of response selection and execution, control networks may lose modular structure as the circuitry collectively converges on a behavioral choice. This interpretation accords with the fact that “choice axes” are encoded within multiple key nodes of frontoparietal decision circuitry (e.g., in macaque LIP: Roitman & Shadlen, 2002; in macaque caudal DLPFC: Mante, Sussillo, Shenoy, & Newsome, 2013; in human DACC and pre-SMA: Minxha, Adolphs, Fusi, Mamelak, & Rutishauser, 2020; see also Okazawa, Hatch, Mancoo, Machens, & Kiani, 2021). This account could be addressed using the enriched experimental design described above, identifying when and where choice coding is emphasized over the course of a trial.

Another unexpected finding was a relatively robust negative relationship between the strength of incongruency coding in left ventral visual cortex and the magnitude of the Stroop effect (Figure 3, B, left panel). One interpretation of this finding is provided by the biased competition framework, as an effect of selective visual attention. Prior work has demonstrated that certain ventral visual regions, those which are strongly tuned to target features, activate as a function of Stroop incongruency (Egner & Hirsch, 2005). In our task, mid-ventral stream areas may have received biasing input, selectively on incongruent trials, which enhanced stimulus-related coding and communication with downstream regions. Using the expanded RSA design sketched above, we might expect that such an effect would be limited to color naming conditions, when selective attention processes would be most prominent.

Perhaps the most surprising result was the robust negative correlation observed between left DLPFC distractor coding and the behavioral Stroop effect (Figure 3, B, right panel). At face, accounting for this finding within the framework of top-down biased competition is difficult. But, given the statistics of our task, in which incongruent trials were frequent and congruent were rare, distractor information could have been used to facilitate performance. An association between distractor features and incongruency could have been learned and used to influence response selection — for example, by retrieving and implementing a stimulus-appropriate attentional setting (Bugg & Crump, 2012; Melara & Algom, 2003). In fact, subjects clearly do exploit these associations, as indicated by reduced Stroop effects for stimuli that are “mostly incongruent,” also known as “item-specific proportion congruency” effects (ISPC; Bugg, Jacoby, & Chanani, 2011); see also Crump & Milliken, 2009). The prediction that distractor coding might reflect ISPC effects could be tested by varying ISPC levels across different stimuli. For stimuli in which the specific color or word is not predictive of congruency, the relationship between DLPFC distractor coding and improved Stroop performance should not be present.

Finally, the present study sets the stage for using RSA to test the dual-mechanisms framework of cognitive control (Braver, 2012). This framework explains much within and between-individual variability in cognitive control function by the existence of two operational “modes” of cognitive control: proactive and reactive. These modes are proposed to have dissociable signature neural coding schemes. Proactive control should rely heavily on goal-relevant coding schemes that originate in LPFC prior to target-stimulus onset as abstract rule or context coding, but which may morph into target coding post-stimulus onset. In contrast, reactive control should rely on an incongruency-based coding-scheme (including coding of whichever task dimensions are predictive of incongruency), originating post-target onset, with potential loci in DMFC or subcortical structures (Chiu, Jiang, & Egner, 2017; Jiang, Brashier, & Egner, 2015). As suggested here, it may be possible to measure correlates of these neural coding schemes via RSA. Experimental factors that encourage subjects to adopt one mode over another (e.g., strategy training, expectancy of difficulty) should correspondingly shift frontoparietal coding schemes along these proactive and reactive dimensions. Further, their behavioral relevance should predictably change, as well: for example, in task contexts in which a proactive control mode is theoretically maladaptive, subjects with stronger proactive coding should perform worse. Thus, the dual-mechanisms framework suggests a broad range of hypotheses amenable to testing with RSA methods (see, e.g., Hall-McMaster, Muhle-Karbe, Myers, & Stokes, 2019). Such hypotheses can be addressed in the broader Dual Mechanisms of Cognitive Control dataset (Braver, Kizhner, Tang, Freund, & Etzel, 2020), of which the data used here are a small subset.

## Acknowledgements

Funding support for this work was provided to TSB through NIH grant R37 MH066078. We thank all former and current team members of the Dual Mechanisms of Cognitive Control Project for their efforts, Jo Etzel for general methodological wisdom, Atsushi Kikimoto for useful thoughts on our RSA models, the Cognitive Control and Psychopathology Laboratory for support and suggestions, and also other WUSTL colleagues in the Bugg, Kool, and Zachs laboratories. Mensh and Kording (2017) was a useful resource for organizing an initial draft of this manuscript. Computations were performed in part using the facilities of the Washington University Center for High Performance Computing, which were partially funded by NIH grants 1S10RR022984-01A1 and 1S10OD018091-01. Surface images were prepared with *Connectome Workbench* (Marcus et al., 2011). The original copy of this manuscript was drafted with *papaja* (Aust & Barth, 2020) and *knitr* (Xie, 2015).

## Extended Data

**Figure 1-1:**
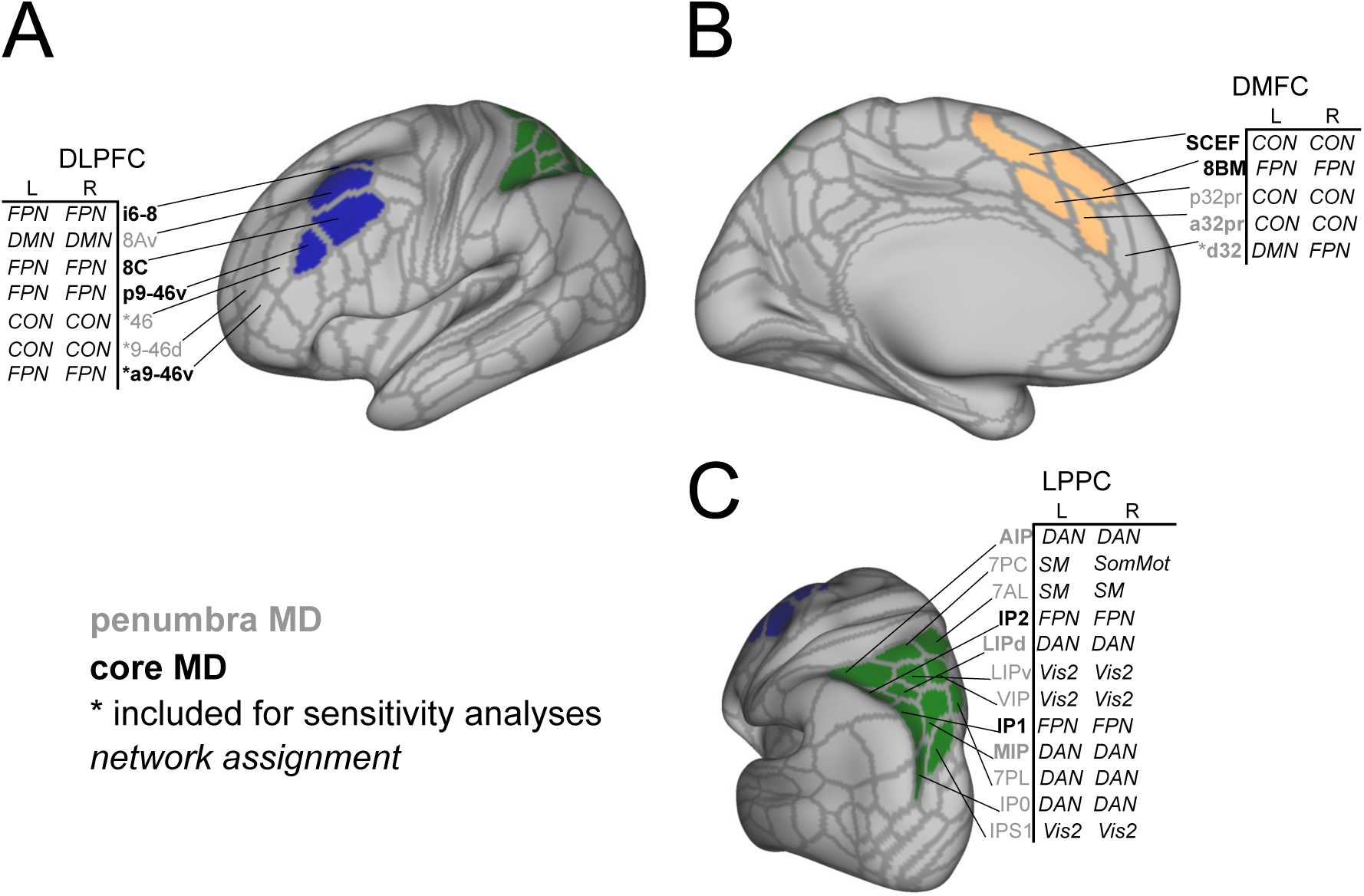
Regions of interest, or “superparcels,” for primary analyses. Colored parcels were included in these superparcels. For DLPFC and DMFC, we additionally conducted sensitivity analyses with more inclusive definitions (i.e., including parcels with asterisked labels). Network assignments were provided in Ji et al. (2019). Parcel names that are in bold were identified as likely being situated over nodes of the “multiple demand” network (Assem, et al. 2020), a collection of regions that is commonly recruited by a variety of demanding tasks. Of these, those in grey belonged to the penumbra (recruited somewhat task-selectively) and those in black belonged to the core (robustly recruited regardless of task). DLPFC, dorsolateral prefrontal cortex; DMFC, dorsomedial frontal cortex; LPPC, lateral posterior parietal cortex; MD, multiple demand; FPN, frontoparietal network; CON, cinguloopercular network; DMN, default mode network; DAN, dorsal attention network; SM, somatomotor network; Vis2, 2nd visual network.

**Figure 1-2:**
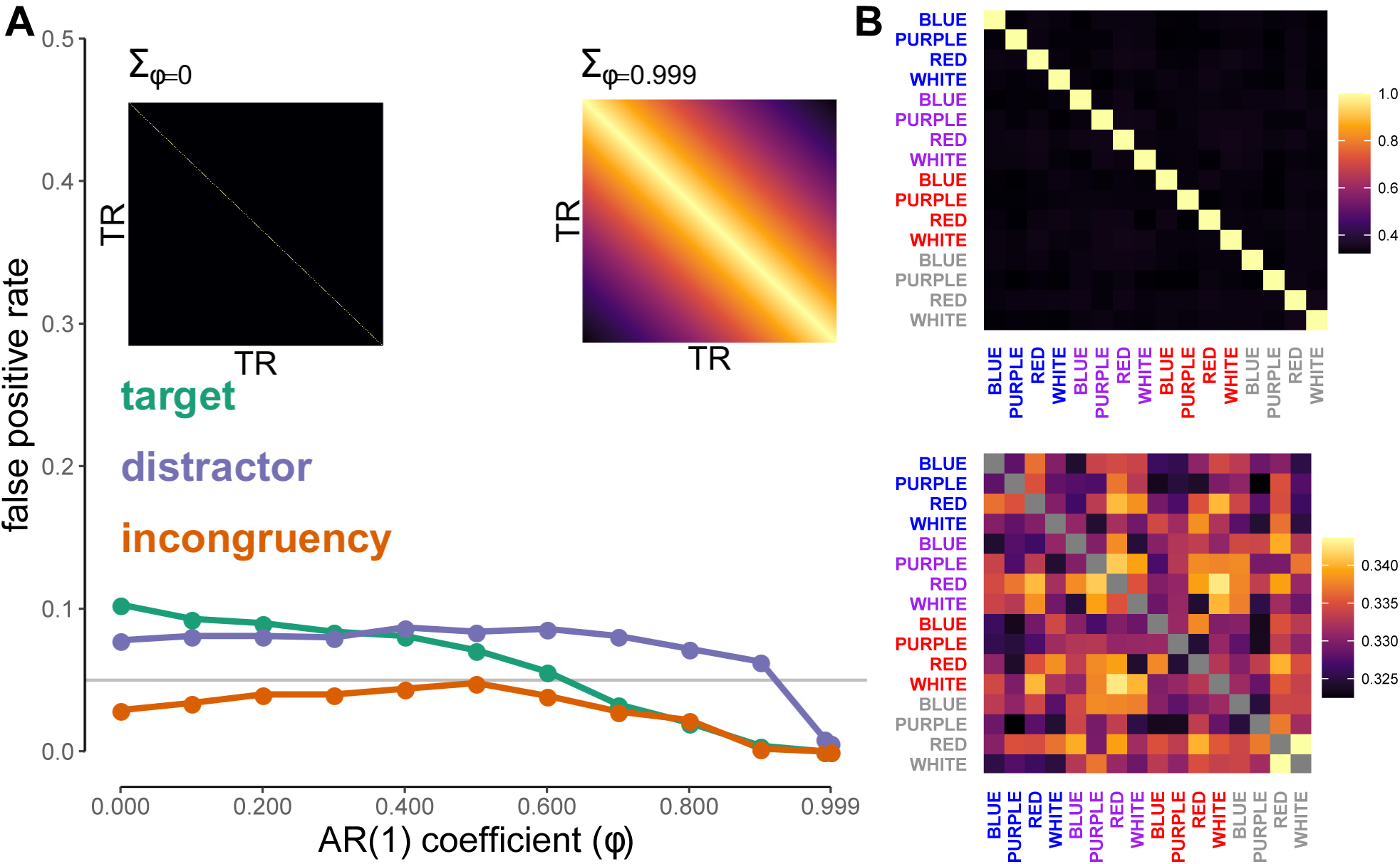
Simulation to diagnose the presence and extent of design bias. **A**, Estimated false positive rate of each RSA model across a range of autocorrelation strengths. We simulated time-series of pure noise and regressed these onto each subjects’ design matrix used in the analysis. Time-series were drawn **Y***_t×v_ ∼ N_t_*(**0**, **Σ***_t×t_*), for *t* in 1, …, 1080 TRs and *v* in 1, …, 100 voxels. Multiple **Σ***_t×t_*s were generated to have AR(1) structure across a range of AR coefficients, *ϕ* in 0, 0.1, 0.2, …, 0.9, 0.99, 0.999. For each *ϕ*, 10,000 simulations were conducted. For each simulation, GLM coefficient estimates from these regressions were submitted to RSA as outlined in *Estimation of Coding Strength β*, and the rate of significantly greater-than-zero RSA model fits was tallied as the false positive rate. At worst (low levels of autocorrelation), models were weakly biased. Note that an empirically-derived false positive rate, obtained from fitting RSA models to patterns of “activation” within ventricles, additionally indicated a false positive rate of no worse than than 0.07 (see Method). **B**, the average RSA correlation matrix from the “worst-case” simulation (*ϕ* = 0). In the top panel, the color-scale ranges from ~0.3 to 1. As the off-diagonal appears homogeneous, the scale of the bias was quite small. In the bottom panel, the diagonal (all 1s) is masked so the color-scale shows the bias structure. No “representational” structure is readily apparent (*cf*., Cai et al., 2020).

**Figure 2-1:**
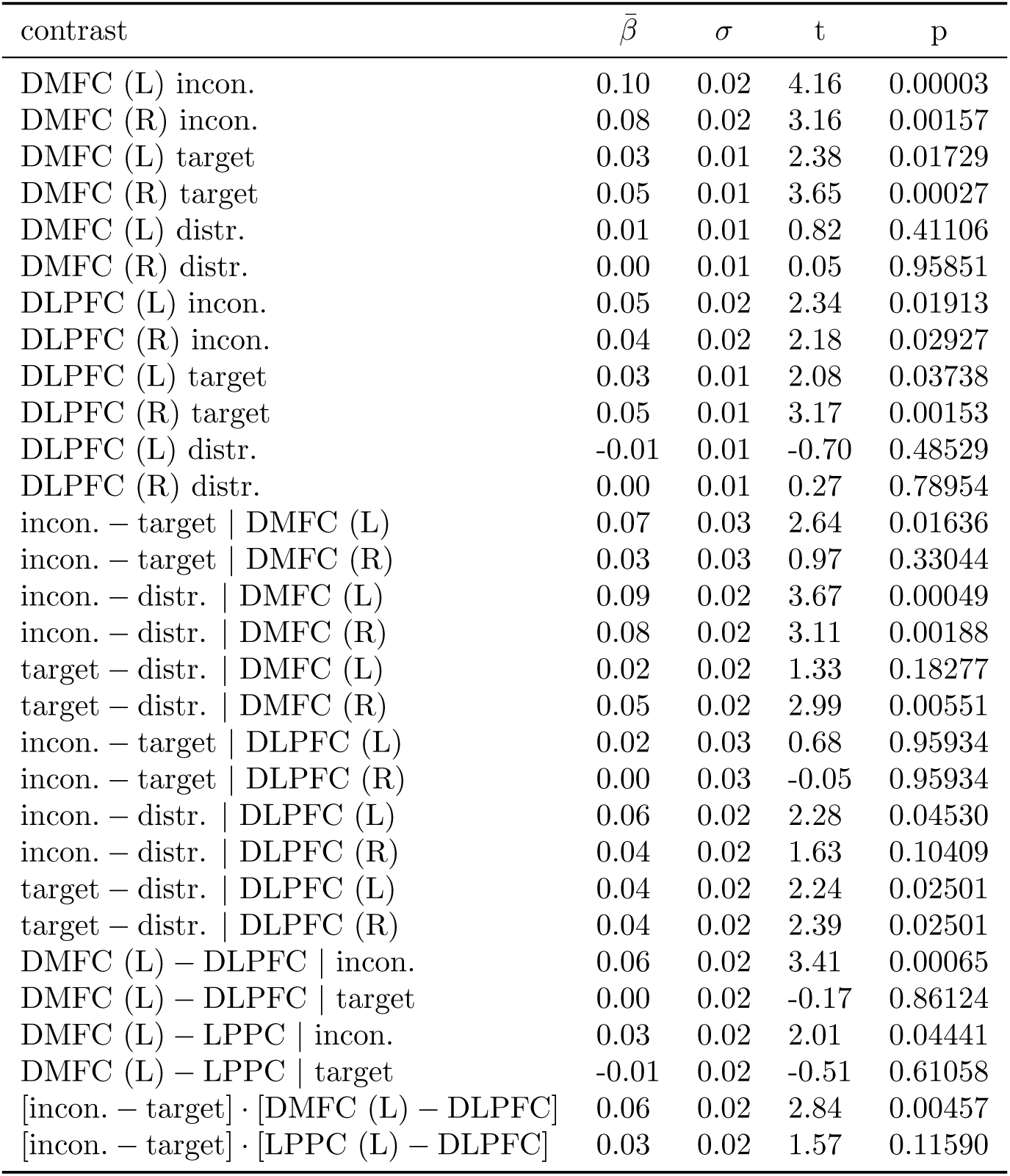
Parameter estimates from a sensitivity test of the group-level RSA, which used alternative (more inclusive) definitions of DLPFC and DMFC (see Figure 1-1). This table contains statistics analogous to Table 2. 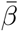 indicates mean RSA model fit over subjects. Unless noted, contrasts were collapsed across hemisphere. For contrasts performed separately in each hemisphere, p-values were FDR-adjusted for multiple comparisons. Incon., incongruency; distr., distractor.

**Figure 2-2:**
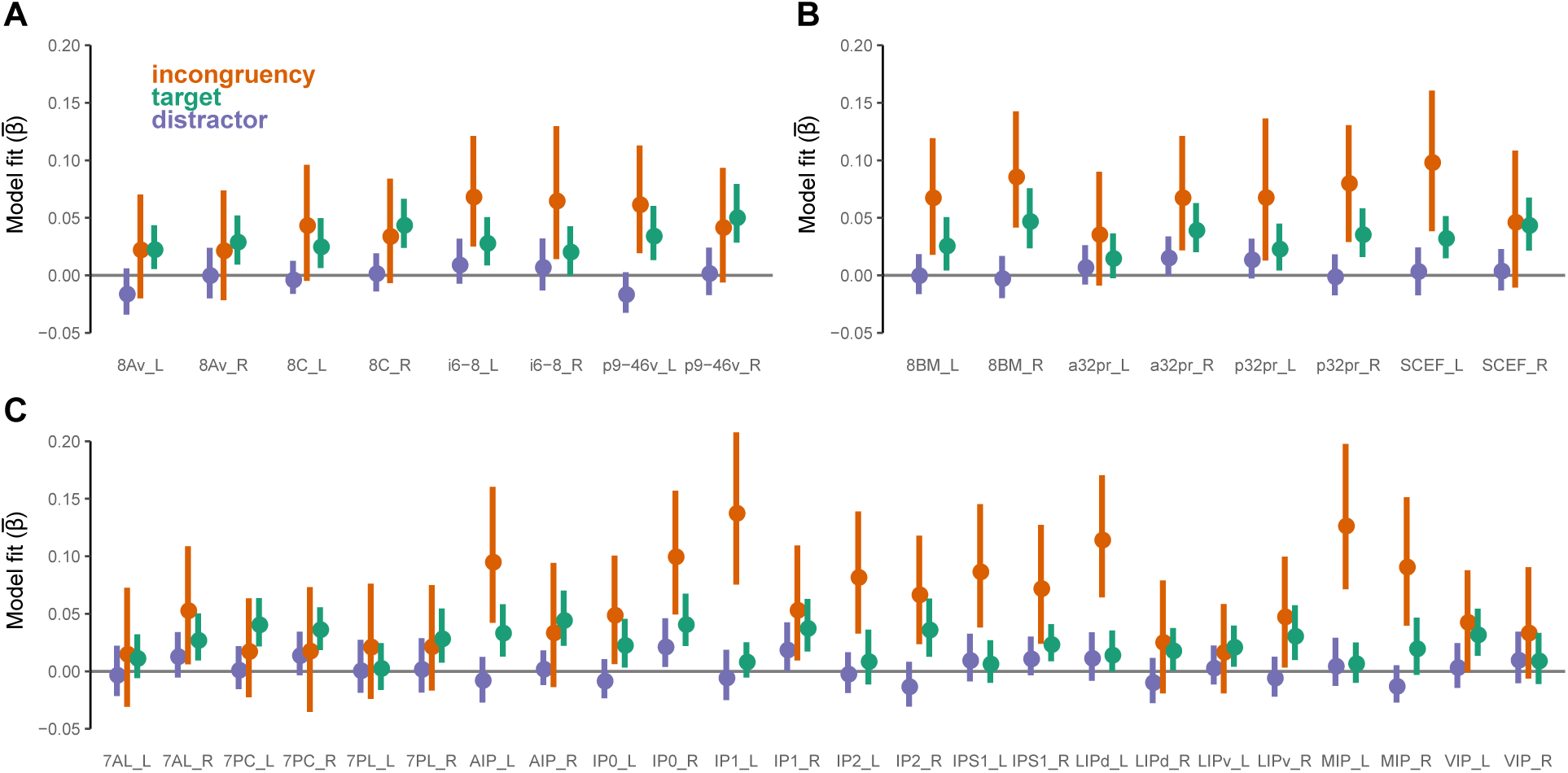
Mean RSA model fits in each MMP parcel within frontoparietal ROIs. (**A**) DLPFC, (**B**) DMFC, (**C**) LPPC. Hemispheres displayed separately. Error-bars represent 95% confidence intervals on between subject variability (estimated via bias-corrected and accelerated bootstrap, 10,000 resamples).

**Figure 2-3:**
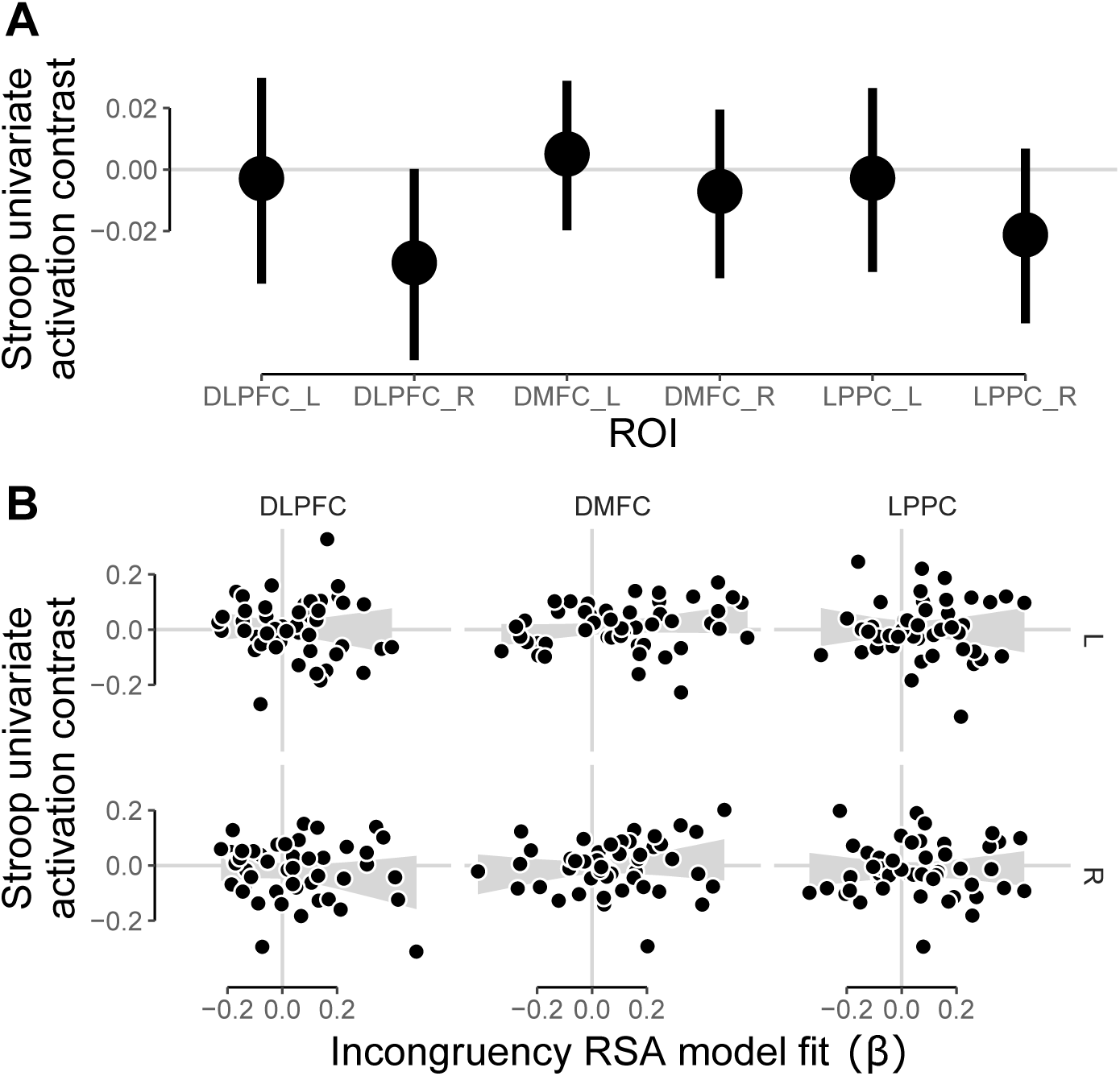
Univariate analysis. **A**, Across-voxel mean levels of activity in response to incongruent versus congruent conditons (“univariate Stroop contrast”) within each frontoparietal ROI. Error-bars represent 95% confidence intervals of between-subject variability (estimated via bootstrap). **B**, Subject-level correlations between the univariate Stroop contrast and RSA incongruency model fit.

**Figure 2-4:**
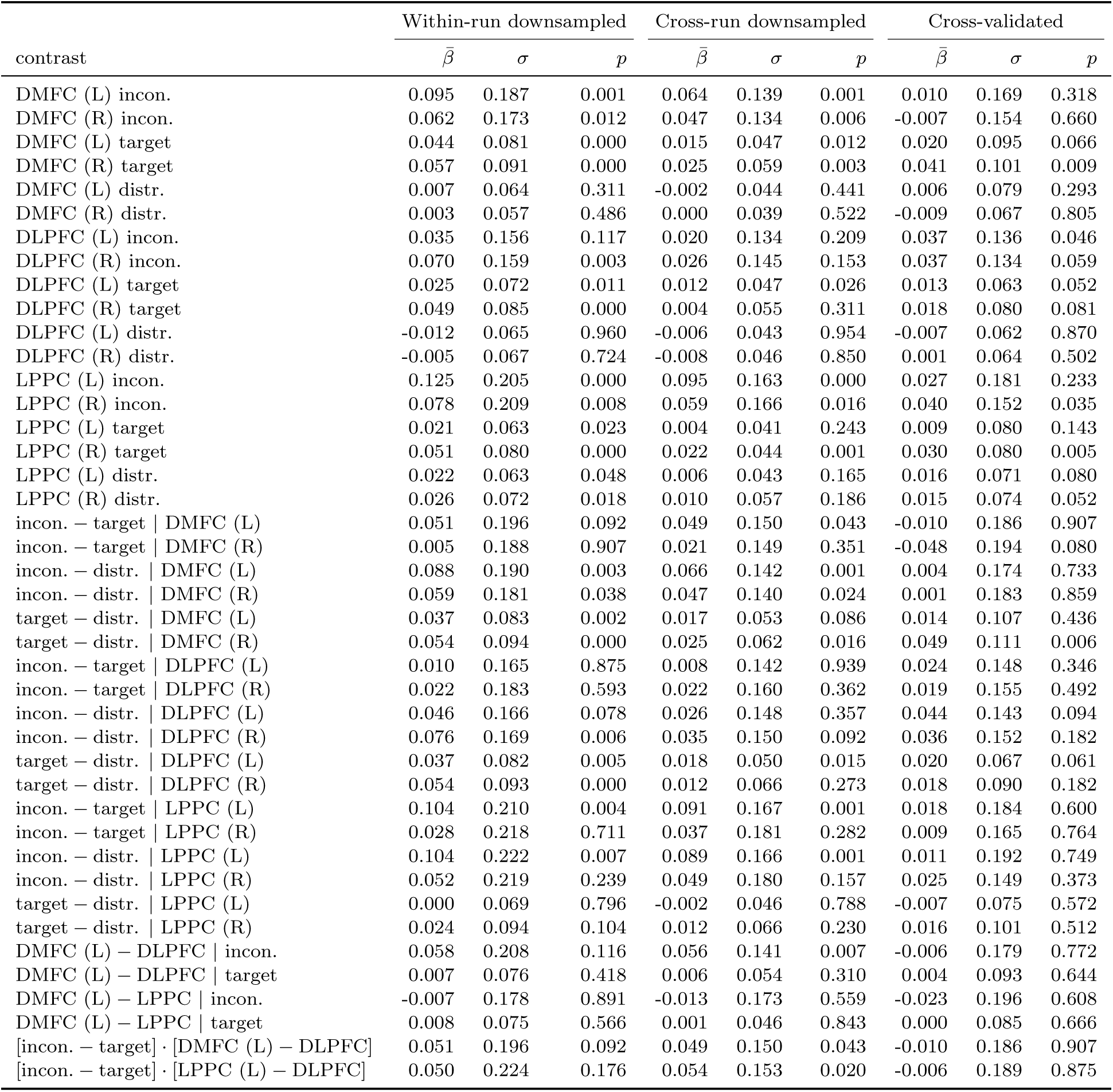
Group-level results from alternative RSA techniques. “Within-run downsampled” columns correspond to results from the downsampling analysis, while “Cross-run downsampled” and “Cross-validated” columns correspond to results from between-run RSA (see relevant Method sections). Following Nili et. al (2014), one-sample contrasts are sign-rank tested against a one-tailed hypothesis, while model comparisons are two-tailed. P-values uncorrected.

**Figure 2-5:**
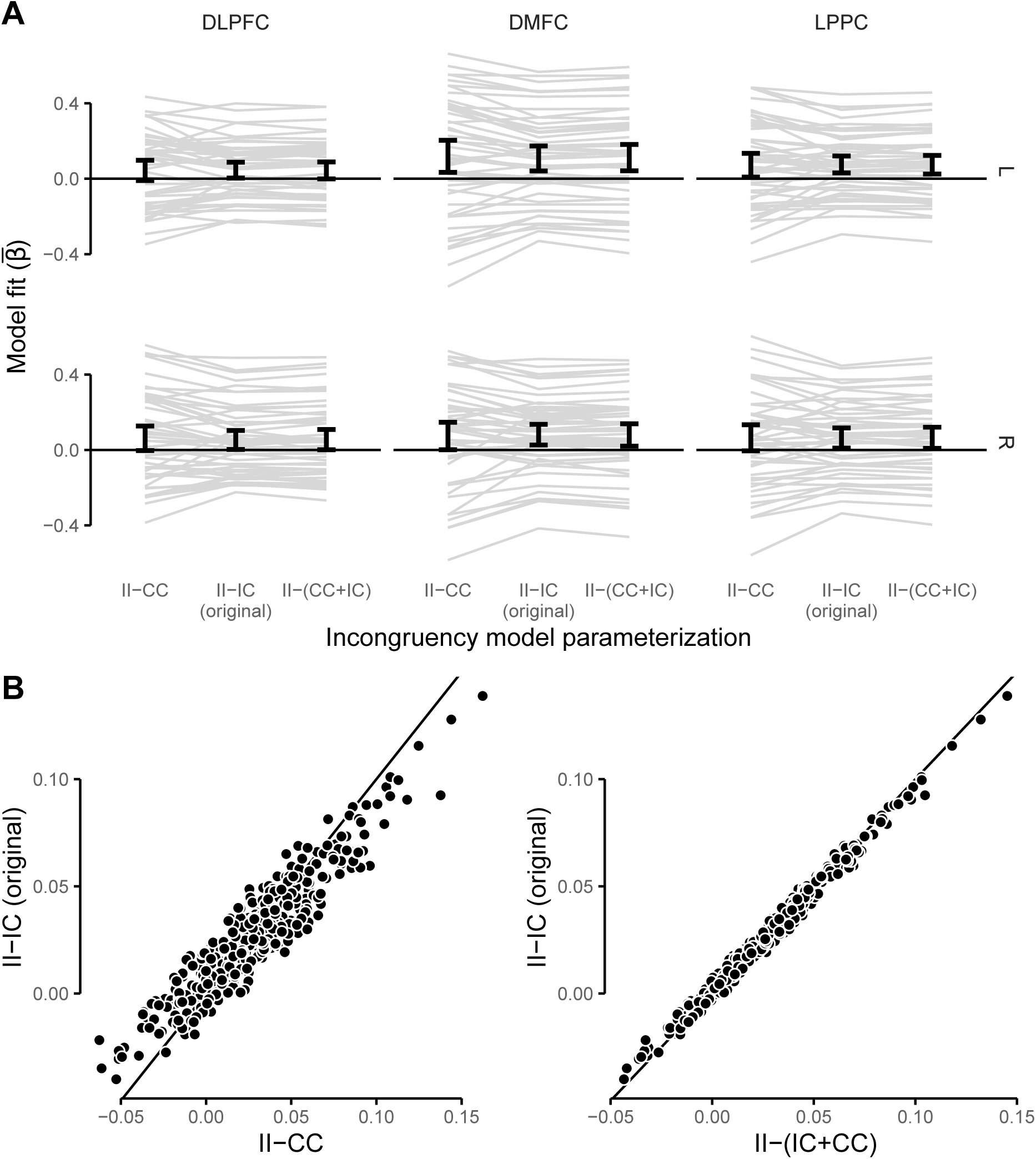
Comparison of results from alternative RSA incongruency models to original parameterization. Two alternative incongruency models were fitted, and model fits (*β*) were compared to our original parameterization of incongruency coding. See *Alternative RSA Incongruency Models* section of Methods for description of alternative models. Note that the *II − CC* parameterization is expected to yield higher variance fits than the other two, due to excluding substantially more data per subject. **A**, Incongruency model fits for each parameterization in each of our ROIs. Error bars show bootstrapped 95% CI. Each line is a subject. Model fits were highly similar across parameterizations. **B**, Incongruency model fits for each parameterization in each MMP parcel (each point is a parcel). Model fits were highly similar across parameterizations.

**Figure 3-1:**
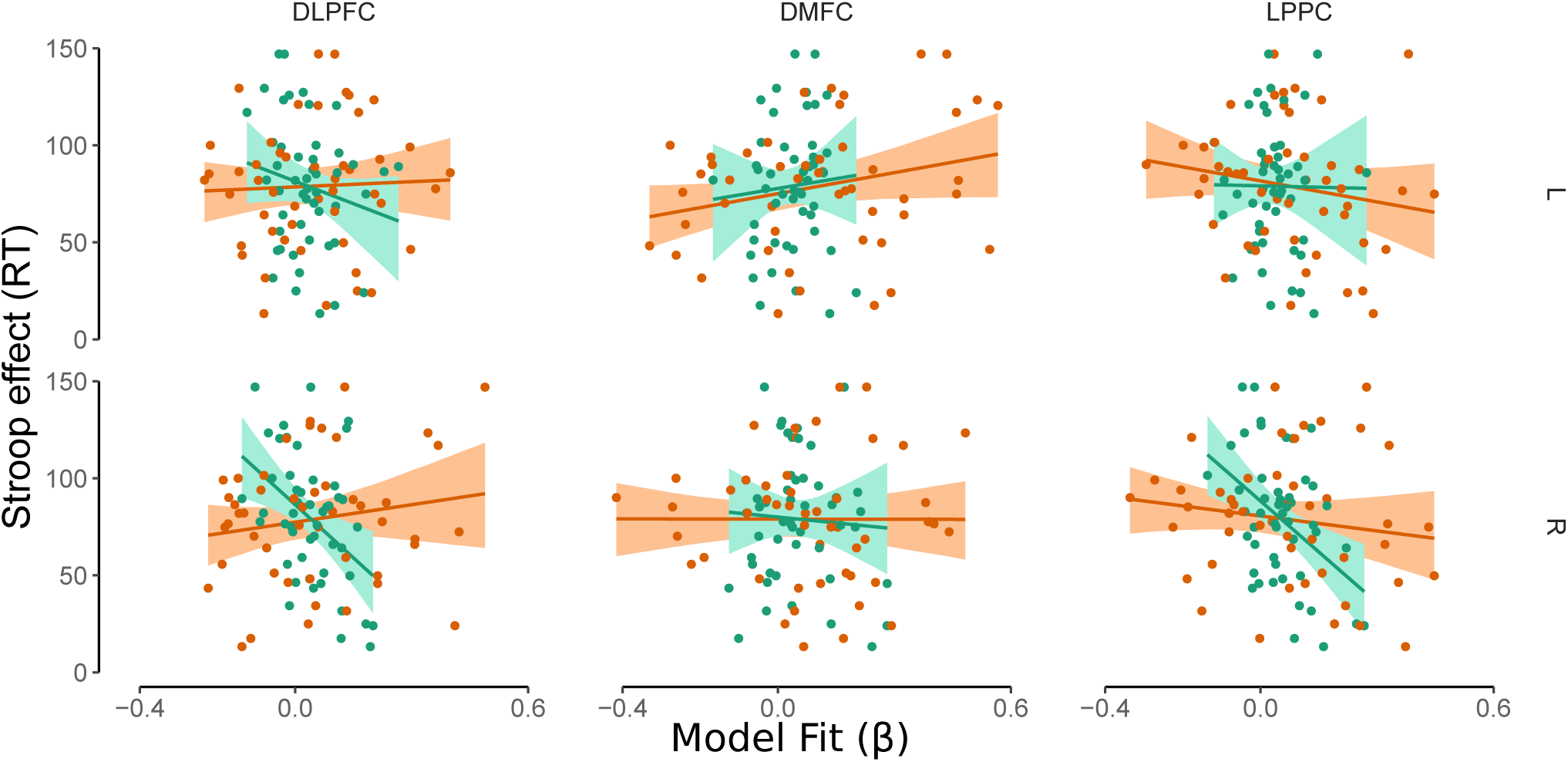
Associations between subjects’ coding strength (*β*) and Stroop interference effect (response time) separately displayed within each hemisphere, ROI, and coding model (target, incongruency). Green, target coding model; orange; incongruency coding model. Confidence bands indicate 95% confidence interval from percentile bootstrap (10,000 resamples).

**Figure 3-2:**
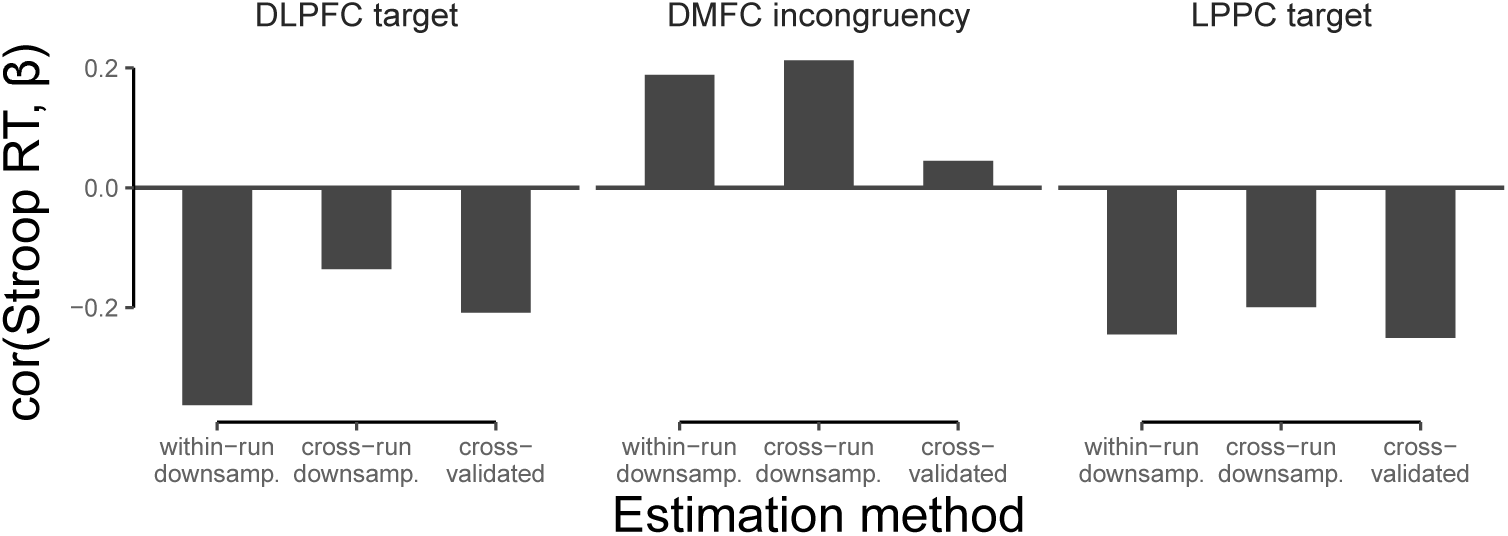
Robustness check of individual-level correlations found in our primary analysis (see Figure 3 and Table 3) to alternative RSA techniques. “Within-run downsampled” corresponds to results from the downsampling analysis, while “Cross-run downsampled” and “Cross-validated” corresponds to the between-run RSA (see relevant Method sections). While attenuation of effect sizes is generally expected when moving to higher variance methods (e.g., downsampling, between-run RSA), critically, the signs of the effects are robust to technique. Notably, however, the large reduction in DMFC incongruency coding correlation with cross-validated RSA, but preservation of the effect with cross-run RSA, matches what is seen at the group level (Figure 2-4, Figure 4-4), and suggests that incongruency information was encoded non-linearly within DMFC activation patterns.

**Figure 3-3:**
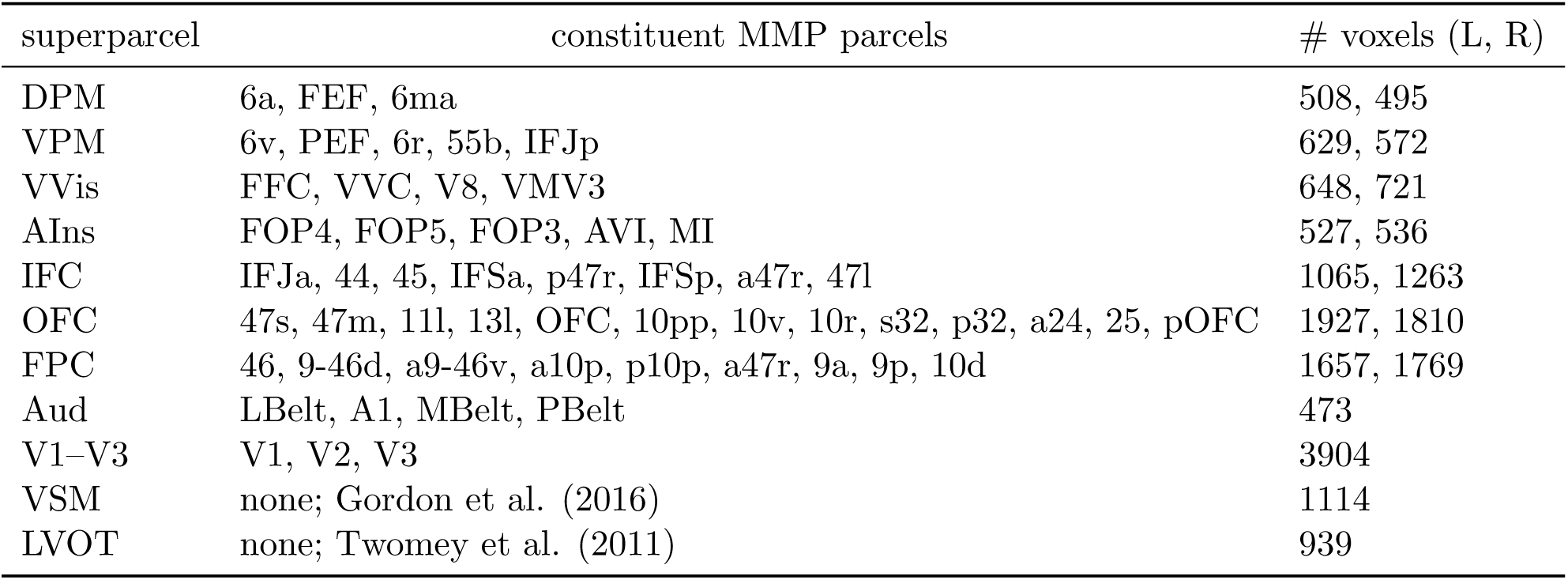
Superparcels defined for model-selection analysis. All superparcels were lateralized (one in each hemisphere) except for relatively early sensory and motor regions (‘Aud,’ ‘V1–V3,’ ‘VSM’). Ventral somatomotor cortex was defined using the Gordon et al. (2016) atlas ‘somato-motor–mouth network,’ as MMP’s somatomotor parcels encompass all of somatomotor strip (i.e., dorsal and ventral areas). Left ventral occipitotemporal cortex (“visual word-form area”) was defined using coordinates from Twomey et al. (2011). DPM, dorsal premotor cortex; VPM, ventral premotor cortex; VVis, ventral visual cortex; AIns, anterior (frontal) insular cortex; IFC, inferior frontal cortex; OFC, orbitofrontal cortex; FPC, frontoparietal cortex; Aud, early auditory cortex; V1–V3, visual cortex, VSM, ventral somatomotor cortex. LVOT, left visual occipitotemporal cortex.

**Figure 4-1:**
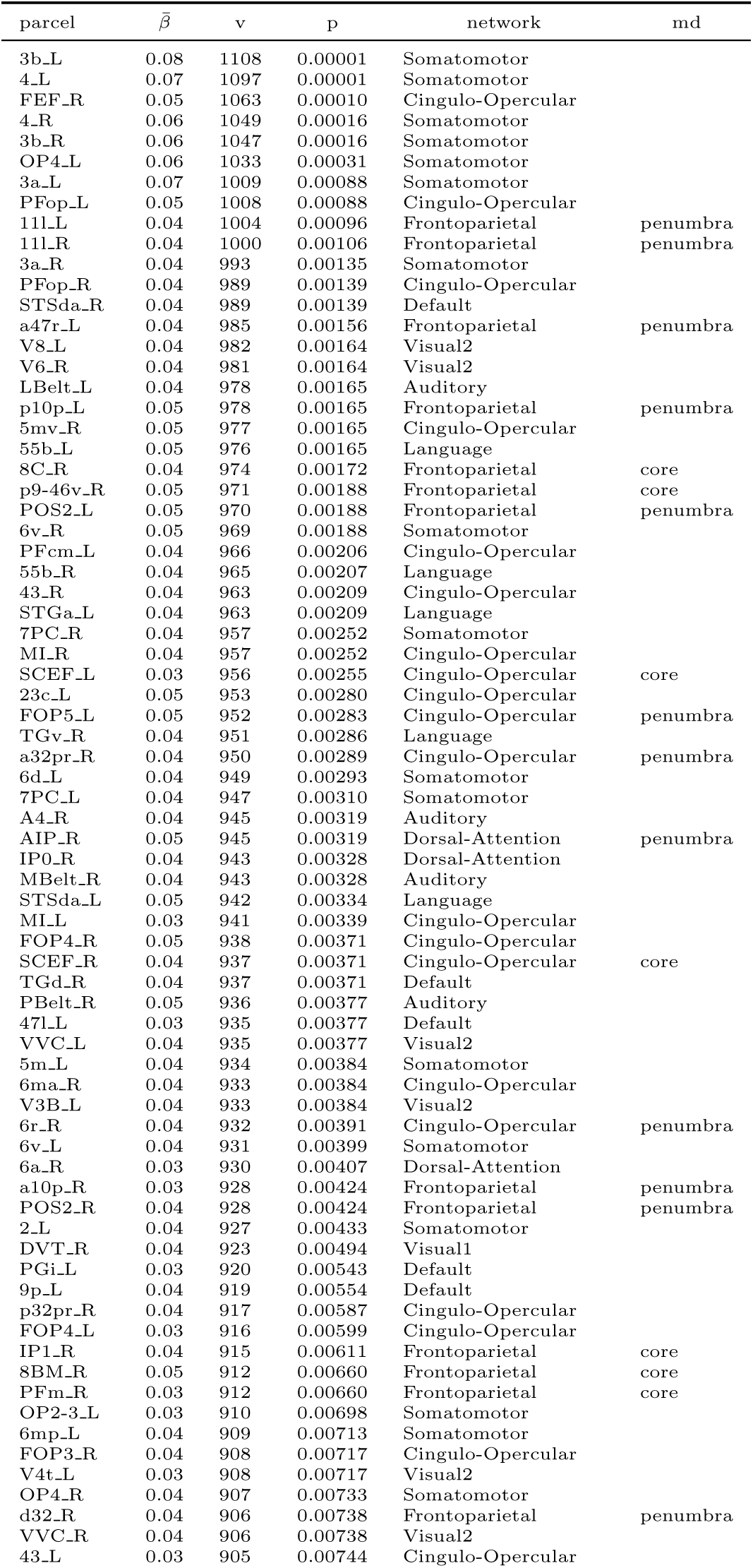
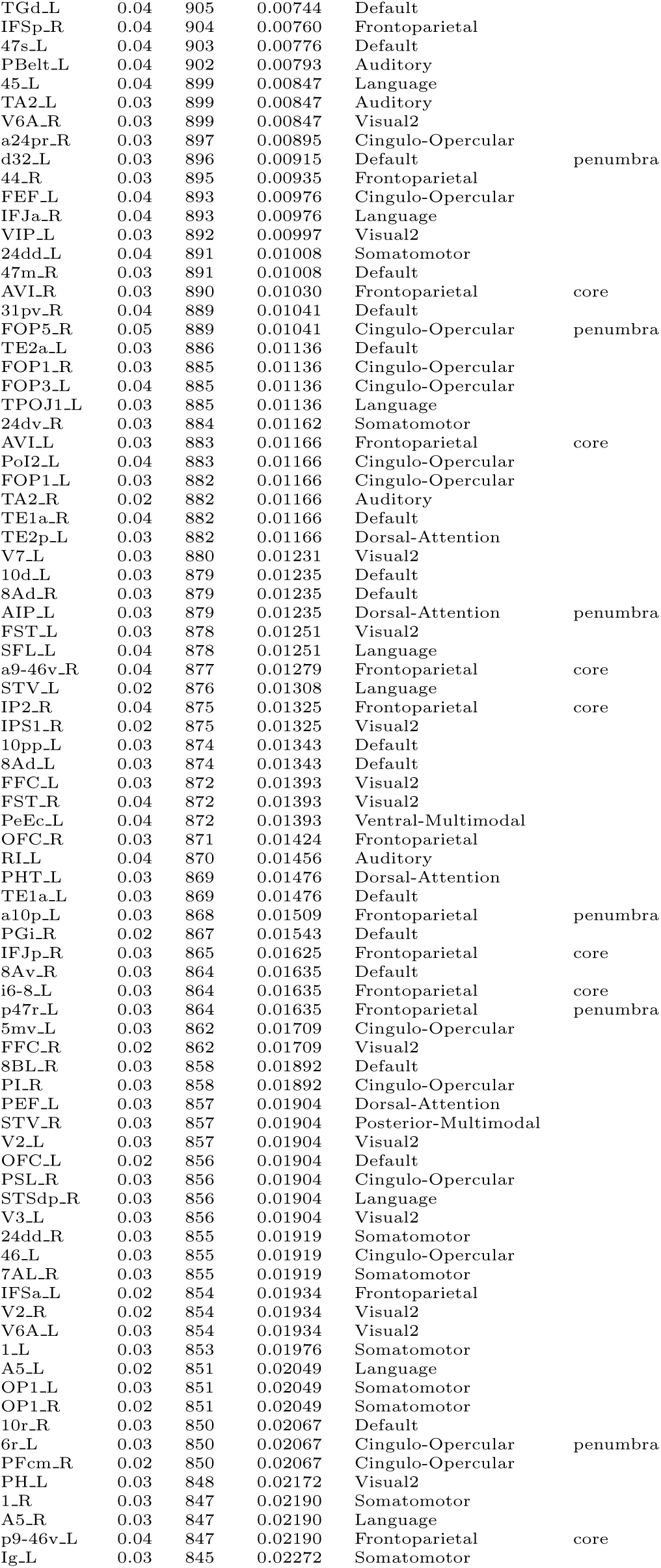
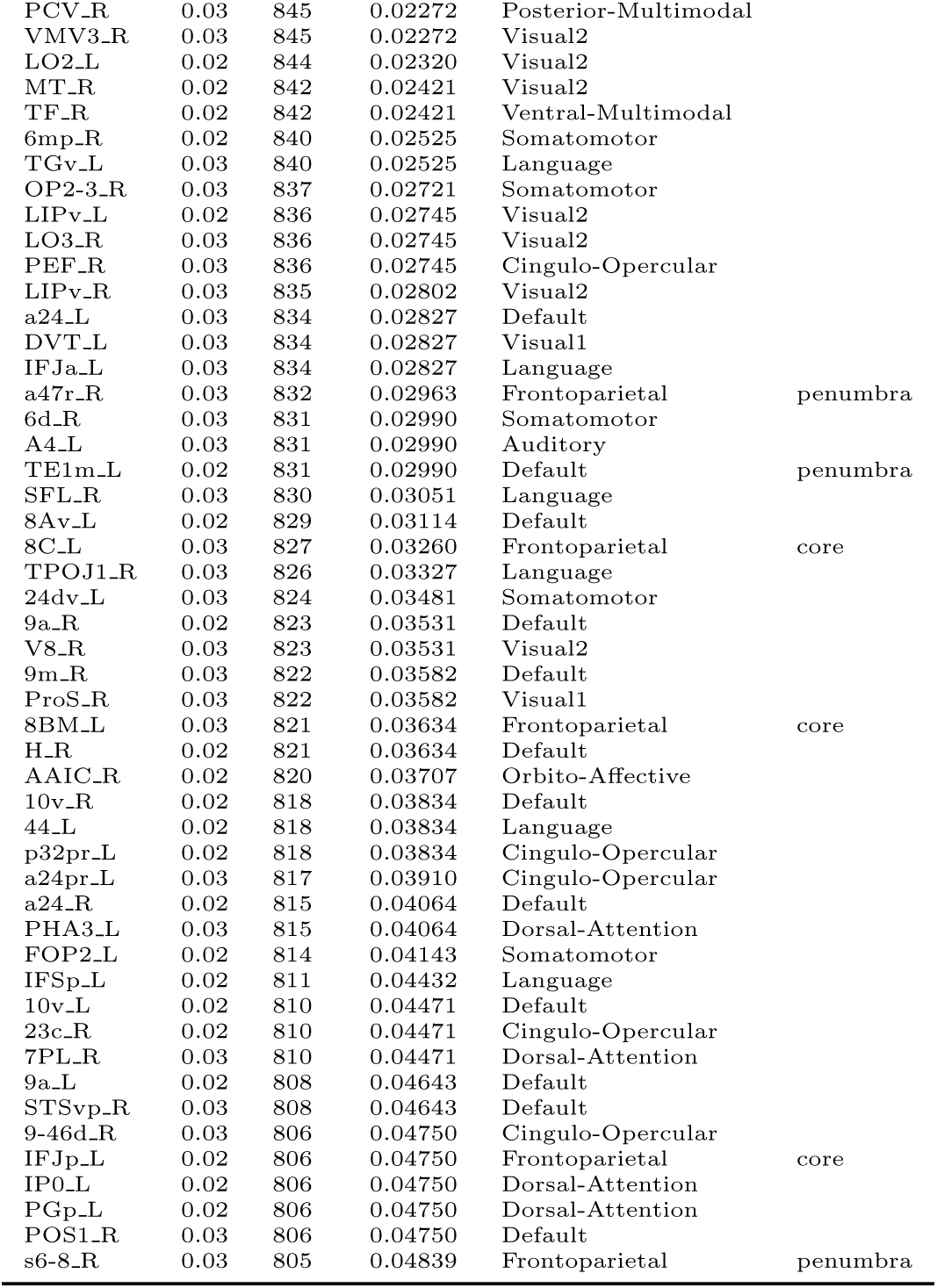
MMP parcels with significant target model fits. P-values are FDR-corrected across all 360 cortical parcels. 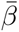, Across-subject mean of RSA model fits. v, Wilcoxon sum of signed-rank test statistic. network, Cole-Anticevic network assignemnt. md, Multiple-Demand assignment (Assem, et al. 2020).

**Figure 4-2:**
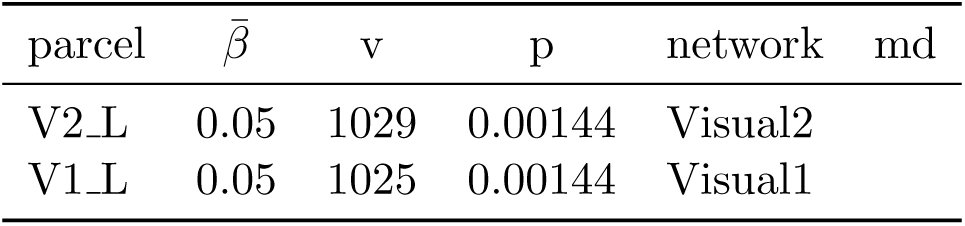
MMP parcels with significant distractor model fits. P-values are FDR-corrected across all 360 cortical parcels. 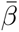, Across-subject mean of RSA model fits. v, Wilcoxon sum of signed-rank test statistic. network, Cole-Anticevic network assignemnt. md, Multiple-Demand assignment (Assem, et al. 2020).

**Figure 4-3:**
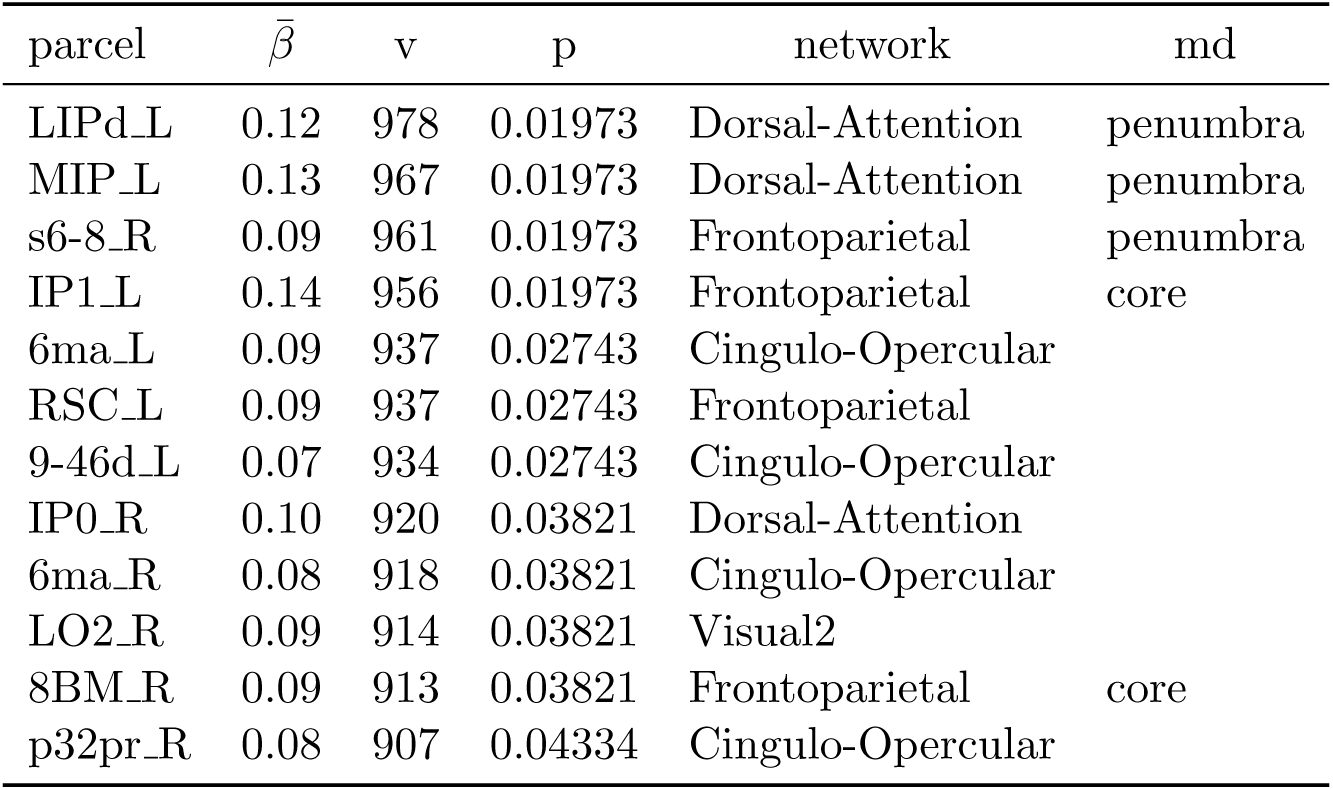
MMP parcels with significant incongruency model fits. P-values are FDR-corrected across all 360 cortical parcels. 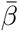, Across-subject mean of RSA model fits. v, Wilcoxon sum of signed-rank test statistic. network, Cole-Anticevic network assignemnt. md, Multiple-Demand assignment (Assem, et al. 2020).

**Figure 4-4:**
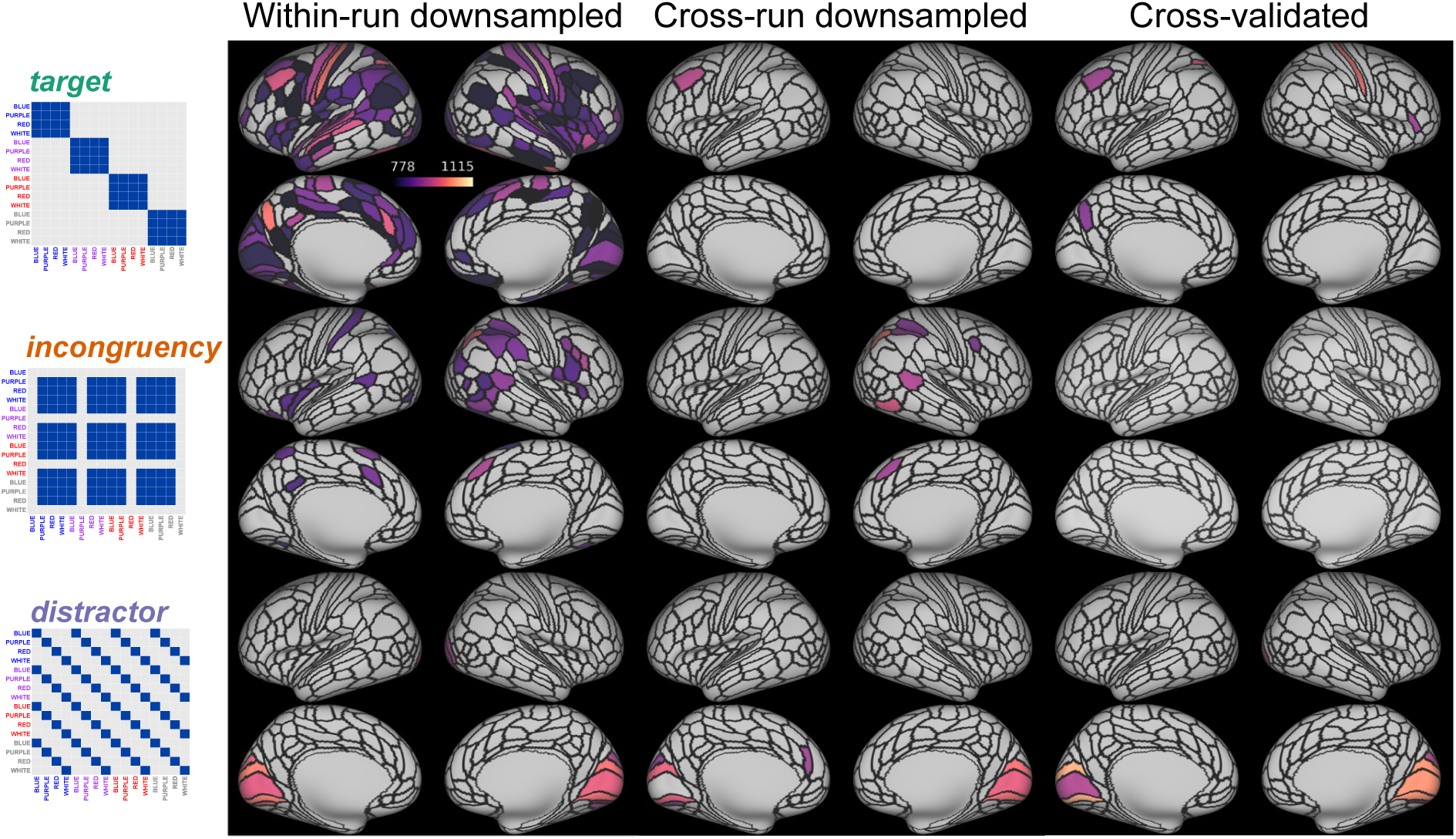
Robustness check of results from exploratory whole-cortex RSA (see Figure 4) to alternative RSA techniques. “Within-run downsampled” corresponds to results from the downsampling analysis, while “Cross-run downsampled” and “Cross-validated” corresponds to between-run RSA (see relevant Method sections). Maps are threhsolded for model fits greater than 0 with significance at FDR-corrected (across all 360 parcels) p-value of 0.05. Note that, in the present design, between-run RSA methods are expected to strongly attenuate effect sizes (see Method). Nevertheless, our core results were robust: in both downsampled and between-run RSA methods, we identified in-congruency coding in parcels within DMFC, target coding in mid-DLPFC, and distractor coding in visual cortex. Notably, the reduction in DMFC incongruency coding with cross-validated RSA, but preservation of the effect with cross-run RSA, matches what is seen at the individual-difference level (Figure 3-2), and suggests that incongruency information was encoded non-linearly within DMFC activation patterns (see also Figure 2, B). Relative to the prewhitening approach we used to handle imbalance of trials across runs, the downsampling approach appears to have increased sensitivity to incongruency coding, in terms of coverage (cf., Figure 4). This could be due to the prewhitening approach (in a separate, preceeding regression) being a somewhat more aggressive denoising strategy. Critically, however, this did not impact our main conclusions: we still observed stronger incongruency coding in DMFC than in DLPFC (see Figure 2-4).

## References

Alink, A., Walther, A., Krugliak, A., Bosch, J. J. F. van den, & Kriegeskorte, N. (2015). Mind the drift - improving sensitivity to fMRI pattern information by accounting for temporal pattern drift. bioRxiv, 032391. https://doi.org/gfsh5f

Allefeld, C., Görgen, K., & Haynes, J.-D. (2016). Valid population inference for information-based imaging: From the second-level t-test to prevalence inference. Neuroimage, 141, 378–392.

Assem, M., Glasser, M. F., Van Essen, D. C., & Duncan, J. (2020). A Domain-General Cognitive Core Defined in Multimodally Parcellated Human Cortex. Cerebral Cortex, 30 (8), 4361–4380. https://doi.org/ghbh24

Aust, F., & Barth, M. (2020). papaja: Create APA manuscripts with R Markdown. Retrieved from https://github.com/crsh/papaja

Baayen, R. H., & Milin, P. (2010). Analyzing reaction times. International Journal of Psychological Research, 3 (2), 12. https://doi.org/gddhjp

Banich, M. T., Smolker, H. R., Snyder, H. R., Lewis-Peacock, J. A., Godinez, D. A., Wager, T. D., & Hankin, B. L. (2019). Turning down the heat: Neural mechanisms of cognitive control for inhibiting task-irrelevant emotional information during adolescence. Neuropsychologia, 125, 93–108.

Barch, D. M., Braver, T. S., Akbudak, E., Conturo, T., Ollinger, J., & Snyder, A. (2001). Anterior cingulate cortex and response conflict: Effects of response modality and processing domain. Cerebral Cortex, 11 (9), 837–848.

Barr, D. J., Levy, R., Scheepers, C., & Tily, H. J. (2013). Random effects structure for confirmatory hypothesis testing: Keep it maximal. Journal of Memory and Language, 68 (3), 255–278. https://doi.org/10.1016/j.jml.2012.11.001

Bates, D., Maechler, M., Bolker, B., & Walker, S. (2014). lme4: Linear mixed-effects models using Eigen and S4. R Package Version, 1 (7), 1–23.

Bhandari, A., Gagne, C., & Badre, D. (2018). Just above Chance: Is It Harder to Decode Information from Prefrontal Cortex Hemodynamic Activity Patterns? Journal of Cognitive Neuroscience, 30 (10), 1473–1498. https://doi.org/10.1162/jocn_a_01291

Botvinick, M. M., Braver, T. S., Barch, D. M., Carter, C. S., & Cohen, J. D. (2001). Conflict Monitoring and Cognitive Control. Psychological Review, 108 (3), 624–652. https://doi.org/10.1037//0033-295X.I08.3.624

Braver, T. S. (2012). The variable nature of cognitive control: A dual mechanisms framework. Trends in Cognitive Sciences, 16 (2), 106–113. https://doi.org/10.1016/j.tics.2011.12.010

Braver, T. S., Kizhner, A., Tang, R., Freund, M. C., & Etzel, J. A. (2020). The Dual Mechanisms of Cognitive Control (DMCC) Project. bioRxiv, 2020.09.18.304402. https://doi.org/ghdrbt

Bugg, Julie M. (2014). Conflict-triggered top-down control: Default mode, last resort, or no such thing? Journal of Experimental Psychology: Learning, Memory, and Cognition, 40 (2), 567.

Bugg, Julie M., & Crump, M. J. C. (2012). In Support of a Distinction between Voluntary and Stimulus-Driven Control: A Review of the Literature on Proportion Congruent Effects. Frontiers in Psychology, 3. https://doi.org/gf39wh

Bugg, Julie M., Jacoby, L. L., & Chanani, S. (2011). Why it is too early to lose control in accounts of item-specific proportion congruency effects. Journal of Experimental Psychology: Human Perception and Performance, 37 (3), 844–859. https://doi.org/d6nd5h

Buschman, T. J., & Miller, E. K. (2007). Top-Down Versus Bottom-Up Control of Attention in the Prefrontal and Posterior Parietal Cortices. Science, 315 (5820), 1860–1862. https://doi.org/dw85gh

Cai, M. B., Schuck, N. W., Pillow, J. W., & Niv, Y. (2019). Representational structure or task structure? Bias in neural representational similarity analysis and a bayesian method for reducing bias. PLoS Computational Biology, 15 (5), e1006299.

Carter, C. S., Macdonald, A. M., Botvinick, M., Ross, L. L., Stenger, V. A., Noll, D., & Cohen, J. D. (2000). Parsing executive processes: Strategic vs. Evaluative functions of the anterior cingulate cortex. Proceedings of the National Academy of Sciences, 97 (4), 1944–1948.

Chiu, Y.-C., Jiang, J., & Egner, T. (2017). The caudate nucleus mediates learning of stimulus– control state associations. The Journal of Neuroscience, 37 (4), 1028–1038. https://doi.org/10.1523/JNEUROSCI.0778-16.2017

Corporation, M., & Weston, S. (2019). doParallel: Foreach parallel adaptor for the ‘parallel’ package. Retrieved from https://CRAN.R-project.org/package=doParallel

Cox, D. D., & Savoy, R. L. (2003). Functional magnetic resonance imaging (fMRI) “brain reading”: Detecting and classifying distributed patterns of fMRI activity in human visual cortex. NeuroImage, 19 (2), 261–270. https://doi.org/d2xbcz

Cox, R. W. (1996). AFNI: Software for Analysis and Visualization of Functional Magnetic Resonance Neuroimages. Computers and Biomedical Research, 29 (3), 162–173. https://doi.org/ctwqf6

Crump, M. J. C., & Milliken, B. (2009). The flexibility of context-specific control: Evidence for context-driven generalization of item-specific control settings. Quarterly Journal of Experimental Psychology, 62 (8), 1523–1532. https://doi.org/10.1080/17470210902752096

De Pisapia, N., & Braver, T. S. (2006). A model of dual control mechanisms through anterior cingulate and prefrontal cortex interactions. Neurocomputing, 69 (10-12), 1322–1326.

Diedrichsen, J. (2006). A spatially unbiased atlas template of the human cerebellum. NeuroImage, 33 (1), 127–138. https://doi.org/dgzq5s

Diedrichsen, J., Berlot, E., Mur, M., Schütt, H. H., & Kriegeskorte, N. (2020). Comparing representational geometries using the unbiased distance correlation. arXiv:2007.02789 [Stat].

Dowle, M., & Srinivasan, A. (2019). Data.table: Extension of ‘data.frame’. Retrieved from https://CRAN.R-project.org/package=data.table

Edelman, S., Grill-Spector, K., Kushnir, T., & Malach, R. (1998). Toward direct visualization of the internal shape representation space by fMRI. Psychobiology, 26 (4), 309–231.

Egner, T., & Hirsch, J. (2005). Cognitive control mechanisms resolve conflict through cortical amplification of task-relevant information. Nature Neuroscience, 8 (12), 1784–1790. https://doi.org/10.1038/nn1594

E-Prime 2.0. (2013). Pittsburgh, PA: Psychology Software Tools, Inc.

Esteban, O., Ciric, R., Finc, K., Blair, R. W., Markiewicz, C. J., Moodie, C. A., … & Gorgolewski, J. J. (2020). Analysis of task-based functional MRI data preprocessed with fMRIPrep. Nature protocols, 1–17.

Esteban, O., Markiewicz, C. J., Blair, R. W., Moodie, C. A., Isik, A. I., Erramuzpe, A., … & Gorgolewski, K. J. (2019). fMRIPrep: a robust preprocessing pipeline for functional MRI. Nature methods, 16 (1), 111–116.

Etzel, J. A., Zacks, J. M., & Braver, T. S. (2013). Searchlight analysis: Promise, pitfalls, and potential. NeuroImage, 78, 261–269. https://doi.org/10.1016/j.neuroimage.2013.03.041

Floden, D., Vallesi, A., & Stuss, D. T. (2010). Task Context and Frontal Lobe Activation in the Stroop Task. Journal of Cognitive Neuroscience, 23 (4), 867–879. https://doi.org/b7wrdt

Freund, M. C., Etzel, J. A., & Braver, T. (2021). Neural coding of cognitive control: The representational similarity analysis approach. Trends in Cognitive Sciences. https://doi.org/10.1016/j.tics.2021.03.011

Friedman, J., Hastie, T., & Tibshirani, R. (2010). Regularization paths for generalized linear models via coordinate descent. Journal of Statistical Software, 33 (1), 1–22. Retrieved from http://www.jstatsoft.org/v33/i01/

Glasser, M. F., Coalson, T. S., Robinson, E. C., Hacker, C. D., Harwell, J., Yacoub, E., … Van Essen, D. C. (2016). A multi-modal parcellation of human cerebral cortex. Nature, 536 (7615), 171–178. https://doi.org/10.1038/nature18933

Glasser, M. F., Sotiropoulos, S. N., Wilson, J. A., Coalson, T. S., Fischl, B., Andersson, J. L., … Jenkinson, M. (2013). The minimal preprocessing pipelines for the Human Connectome Project. NeuroImage, 80, 105–124. https://doi.org/f46nj4

Gonthier, C., Braver, T. S., & Bugg, J. M. (2016). Dissociating proactive and reactive control in the Stroop task. Memory & Cognition, 44 (5), 778–788. https://doi.org/10.3758/s13421-016-0591-1

Gordon, E. M., Laumann, T. O., Adeyemo, B., Huckins, J. F., Kelley, W. M., & Petersen, S. E. (2016). Generation and Evaluation of a Cortical Area Parcellation from Resting-State Correlations. Cerebral Cortex, 26 (1), 288–303. https://doi.org/10.1093/cercor/bhu239

Haines, N., Kvam, P. D., Irving, L. H., Smith, C., Beauchaine, T. P., Pitt, M. A., … Turner, B. (2020). Learning from the Reliability Paradox: How Theoretically Informed Generative Models Can Advance the Social, Behavioral, and Brain Sciences (preprint). PsyArXiv. https://doi.org/10.31234/osf.io/xr7y3

Hall-McMaster, S., Muhle-Karbe, P. S., Myers, N. E., & Stokes, M. G. (2019). Reward boosts neural coding of task rules to optimize cognitive flexibility. Journal of Neuroscience, 39 (43), 8549–8561. https://doi.org/ggcf34

Hastie, T., Tibshirani, R., & Friedman, J. (2009). The Elements of Statistical Learning: Data Mining, Inference, and Prediction, Second Edition (2nd ed.). New York: Springer-Verlag.

Haxby, J. V., Gobbini, M. I., Furey, M. L., & Ishai, A. (2001). Distributed and overlapping representations of faces and objects in ventral temporal cortex. Science; Washington, 293 (5539), 2425–2430. https://doi.org/cffn26

Hothorn, T., Bretz, F., & Westfall, P. (2008). Simultaneous inference in general parametric models. Biometrical Journal, 50 (3), 346–363.

Inkscape Project. (2020). Inkscape (Version 0.92.5). Retrieved from https://inkscape.org

Ji, J. L., Spronk, M., Kulkarni, K., Repovš, G., Anticevic, A., & Cole, M. W. (2019). Mapping the human brain’s cortical-subcortical functional network organization. NeuroImage, 185, 35–57. https://doi.org/ggd8dm

Jiang, J., Brashier, N. M., & Egner, T. (2015). Memory meets control in hippocampal and striatal binding of stimuli, responses, and attentional control states. Journal of Neuroscience, 35 (44), 14885–14895. https://doi.org/10.1523/JNEUROSCI.2957-15.2015

Kane, M. J., & Engle, R. W. (2002). The role of prefrontal cortex in working-memory capacity, executive attention, and general fluid intelligence: An individual-differences perspective. Psychonomic Bulletin & Review, 9 (4), 637–671. https://doi.org/bwh9mt

Kriegeskorte, N. (2008). Representational similarity analysis – connecting the branches of systems neuroscience. Frontiers in Systems Neuroscience, 2 (November), 1–28. https://doi.org/10.3389/neuro.06.004.2008

Kriegeskorte, N., & Kievit, R. A. (2013). Representational geometry: Integrating cognition, computation, and the brain. Trends in Cognitive Sciences, 17 (8), 401–412. https://doi.org/10.1016/j.tics.2013.06.007

Kruskal, J. B. (1964). Multidimensional scaling by optimizing goodness of fit to a nonmetric hypothesis. Psychometrika, 29 (1), 1–27. https://doi.org/dk9pcw

Kurtzer, G. M., Sochat, V., & Bauer, M. W. (2017). Singularity: Scientific containers for mobility of compute. PloS one, 12 (5), e0177459.

Laird, N. M., & Ware, J. H. (1982). Random-effects models for longitudinal data. Biometrics, 963–974.

Lakens, D. (2017). Equivalence Tests. Social Psychological and Personality Science, 8 (4), 355–362. https://doi.org/gbf8nt

Logan, G. D., & Zbrodoff, N. J. (1979). When it helps to be misled: Facilitative effects of increasing the frequency of conflicting stimuli in a stroop-like task. Memory & Cognition, 7 (3), 166–174.

MacDonald, A. W., Cohen, J. D., Stenger, V. A., & Carter, C. S. (2000). Dissociating the role of the dorsolateral prefrontal and anterior cingulate cortex in cognitive control. Science, 288 (5472), 1835–1838. https://doi.org/10.1126/science.288.5472.1835

MacLeod, C. M. (1991). Half a century of research on the Stroop effect: An integrative review. Psychological Bulletin, 109 (2), 163–203. https://doi.org/10.1037/0033-2909.109.2.163

Mair, P., & Wilcox, R. (2020). Robust Statistical Methods in R Using the WRS2 Package. Behavior Research Methods, 52, 464–488.

Mante, V., Sussillo, D., Shenoy, K. V., & Newsome, W. T. (2013). Context-dependent computation by recurrent dynamics in prefrontal cortex. Nature, 503 (7474), 78–84. https://doi.org/f5gdn6

Marcus, D., Harwell, J., Olsen, T., Hodge, M., Glasser, M., Prior, F., … Van Essen, D. (2011). Informatics and data mining tools and strategies for the human connectome project. Frontiers in Neuroinformatics, 5, 4.

Melara, R. D., & Algom, D. (2003). Driven by information: A tectonic theory of stroop effects. Psychological Review, 110 (3), 422.

Mensh, B., & Kording, K. (2017). Ten simple rules for structuring papers. PLOS Computational Biology, 13 (9), e1005619. https://doi.org/ckqp

Microsoft, & Weston, S. (2019). Foreach: Provides foreach looping construct. Retrieved from https://CRAN.R-project.org/package=foreach

Miller, E. K., & Cohen, J. D. (2001). An Integrative Theory of Prefrontal Cortex Function. Annual Review of Neuroscience, 24 (1), 167–202. https://doi.org/10.1146/annurev.neuro.24.1.167

Minxha, J., Adolphs, R., Fusi, S., Mamelak, A. N., & Rutishauser, U. (2020). Flexible recruitment of memory-based choice representations by the human medial frontal cortex. Science, 368 (6498). https://doi.org/gg4btz

Mumford, Jeanette A., Davis, T., & Poldrack, R. A. (2014). The impact of study design on pattern estimation for single-trial multivariate pattern analysis. NeuroImage, 103, 130–138. https://doi.org/f6qvfq

Mumford, Jeanette A., Turner, B. O., Ashby, F. G., & Poldrack, R. A. (2012). Deconvolving BOLD activation in event-related designs for multivoxel pattern classification analyses. Neuroimage, 59 (3), 2636–2643.

Nee, D. E., Wager, T. D., & Jonides, J. (2007). Interference resolution: Insights from a meta-analysis of neuroimaging tasks. Cognitive, Affective, & Behavioral Neuroscience, 7 (1), 1–17.

Niendam, T. A., Laird, A. R., Ray, K. L., Dean, Y. M., Glahn, D. C., & Carter, C. S. (2012). Meta-analytic evidence for a superordinate cognitive control network subserving diverse executive functions. Cognitive, Affective, & Behavioral Neuroscience, 12 (2), 241–268.

Nili, H., Wingfield, C., Walther, A., Su, L., Marslen-Wilson, W., & Kriegeskorte, N. (2014). A Toolbox for Representational Similarity Analysis. PLoS Computational Biology, 10 (4). https://doi.org/10.1371/journal.pcbi.1003553

Okazawa, G., Hatch, C. E., Mancoo, A., Machens, C. K., & Kiani, R. (2021). The geometry of the representation of decision variable and stimulus difficulty in the parietal cortex. bioRxiv, 2021–2001.

Oksanen, J., Blanchet, F. G., Friendly, M., Kindt, R., Legendre, P., McGlinn, D., … Wagner, H. (2019). Vegan: Community ecology package. Retrieved from https://CRAN.R-project.org/package=vegan

Petersen, S. E., & Dubis, J. W. (2012). The mixed block/event-related design. Neuroimage, 62 (2), 1177–1184.

Petrides, M., & Pandya, D. (1999). Dorsolateral prefrontal cortex: Comparative cytoarchitectonic analysis in the human and the macaque brain and corticocortical connection patterns. European Journal of Neuroscience, 11 (3), 1011–1036.

Pinheiro, J., Bates, D., DebRoy, S., Sarkar, D., & R Core Team. (2019). nlme: Linear and nonlinear mixed effects models. Retrieved from https://CRAN.R-project.org/package=nlme

Posner, M. I., & Snyder, C. R. R. (1975). Attention and Cognitive Control. In *In* R. Solso *(Ed.)* Information processing and cognition: The Loyola Symposium (pp. 669–682).

R Core Team. (2019). R: A language and environment for statistical computing. Vienna, Austria: R Foundation for Statistical Computing. Retrieved from https://www.R-project.org/

Ridderinkhof, K. R., Ullsperger, M., Crone, E. A., & Nieuwenhuis, S. (2004). The Role of the Medial Frontal Cortex in Cognitive Control. Science, 306 (5695), 443–447. https://doi.org/dzbqvx

Rigotti, M., Barak, O., Warden, M. R., Wang, X. J., Daw, N. D., Miller, E. K., & Fusi, S. (2013). The importance of mixed selectivity in complex cognitive tasks. Nature, 497 (7451), 585–590. https://doi.org/10.1038/nature12160

Roitman, J. D., & Shadlen, M. N. (2002). Response of Neurons in the Lateral Intraparietal Area during a Combined Visual Discrimination Reaction Time Task. The Journal of Neuroscience, 22 (21), 9475–9489. https://doi.org/10.1523/JNEUROSCI.22-21-09475.2002

Schuirmann, D. J. (1987). A comparison of the Two One-Sided Tests Procedure and the Power Approach for assessing the equivalence of average bioavailability. Journal of Pharmacokinetics and Biopharmaceutics, 15 (6), 657–680. https://doi.org/bsdb55

Shenhav, A., Botvinick, M. M., & Cohen, J. D. (2013). The Expected Value of Control: An Integrative Theory of Anterior Cingulate Cortex Function. Neuron, 79 (2), 217–240. https://doi.org/10.1016/j.neuron.2013.07.007

Simmons, J. P., Nelson, L. D., & Simonsohn, U. (2011). False-Positive Psychology: Undisclosed Flexibility in Data Collection and Analysis Allows Presenting Anything as Significant. Psychological Science, 22 (11), 1359–1366. https://doi.org/bxbw3c

Smith, E. H., Horga, G., Yates, M. J., Mikell, C. B., Banks, G. P., Pathak, Y. J., … others. (2019). Widespread temporal coding of cognitive control in the human prefrontal cortex. Nature Neuroscience, 22 (11), 1883–1891.

Stokes, M. G., Kusunoki, M., Sigala, N., Nili, H., Gaffan, D., & Duncan, J. (2013). Dynamic coding for cognitive control in prefrontal cortex. Neuron, 78 (2), 364–375. https://doi.org/10.1016/j.neuron.2013.01.039

Stroop, J. R. (1935). Studies of interference in serial verbal reactions. Journal of Experimental Psychology, 18 (6), 643–662. https://doi.org/10.1037/h0054651

Twomey, T., Kawabata Duncan, K. J., Price, C. J., & Devlin, J. T. (2011). Top-down modulation of ventral occipito-temporal responses during visual word recognition. NeuroImage, 55 (3), 1242–1251. https://doi.org/b2dccr

Walther, A., Nili, H., Ejaz, N., Alink, A., Kriegeskorte, N., & Diedrichsen, J. (2016). Reliability of dissimilarity measures for multi-voxel pattern analysis. Neuroimage, 137, 188–200.

Wickham, H. (2016). ggplot2: Elegant graphics for data analysis. Springer-Verlag New York. Retrieved from https://ggplot2.tidyverse.org

Wickham, H., François, R., Henry, L., & Müller, K. (2020). Dplyr: A grammar of data manipulation. Retrieved from https://CRAN.R-project.org/package=dplyr

Xie, Y. (2015). Dynamic documents with R and knitr (2nd ed.). Boca Raton, Florida: Chapman; Hall/CRC. Retrieved from https://yihui.name/knitr/

